# Context-dependent transcriptional remodeling of TADs during differentiation

**DOI:** 10.1101/2022.07.01.498405

**Authors:** Sanjay Chahar, Yousra Ben Zouari, Hossein Salari, Anne M Molitor, Dominique Kobi, Manon Maroquenne, Cathie Erb, Audrey Mossler, Nezih Karasu, Daniel Jost, Tom Sexton

## Abstract

Metazoan chromosomes are organized into discrete domains (TADs), believed to contribute to the regulation of transcriptional programs. Despite extensive correlation between TAD organization and gene activity, a direct mechanistic link is unclear, with perturbation studies often showing little effect. To follow TAD dynamics during development, we used Capture Hi-C to interrogate the TADs around key differentially expressed genes during mouse thymocyte maturation, uncovering specific remodeling events. Notably, one TAD boundary was broadened to accommodate RNA polymerase elongation past the border, and sub-domains were formed around some activated genes without changes in CTCF binding. The ectopic induction of one gene was sufficient to recapitulate microdomain formation in embryonic stem cells, providing strong evidence that transcription can directly remodel chromatin structure. These results suggest that transcriptional processes drive complex, but non-universal, chromosome folding patterns that can be important in certain genomic contexts.

## Introduction

Eukaryotic genomes are spatially highly organized, permitting access of transcriptional machinery to the appropriate loci despite extensive compaction for containment within the nucleus (Sexton and Cavalli, 2015). A key architectural feature of metazoan chromosomes is their organization into autonomously folded domains, termed topologically associated domains (TADs), whose arrangement correlates very well with functional demarcation of chromatin regions according to gene expression, epigenetic marks and replication timing (Dixon et al., 2012; Sexton et al., 2012; Pope et al., 2014). TADs have been proposed to ensure appropriate gene expression by limiting the functional range of distal transcriptional enhancers, evidenced by pathologies due to ectopic gene activation on loss of TAD borders (Lupiáñez et al., 2015). TADs may conversely facilitate enhancer activity on their cognate genes by limiting their effective search space within a domain (Zuin et al., 2022). However, experimental perturbations causing acute, extensive loss of TADs have only modest effects on the transcriptome (Nora et al., 2017; Rao et al., 2017; Schwarzer et al., 2017), suggesting that chromatin topology is not necessarily instructive in gene regulation. Alternatively, by stabilizing particular configurations more or less favorable for transcriptional firing, TADs could serve to reduce transcriptional noise and cell-to-cell variability (Ren et al., 2017). Metazoan TAD borders are highly enriched for binding of the insulator protein CCCTC-binding factor (CTCF) (Dixon et al., 2012), with the curious feature that flanking TAD borders predominantly comprise CTCF motifs in convergent orientation (Rao et al., 2014). A popular model for TAD formation is loop extrusion, whereby the ring-like cohesin complex binds and translocates bi-directionally along chromatin, bringing linearly distal regions near to one another. The “collision” of convergent CTCF-bound sites with cohesins is proposed to stall loop extrusion, thus creating metastable interactions between TAD boundaries (Davidson et al., 2019; Fudenberg et al., 2016; Li et al., 2020; Sanborn et al., 2015). Live imaging experiments have visualized such predicted interactions, although they are relatively infrequent and transient (Gabriele et al., 2022). Inversion of CTCF motifs disrupts chromatin interactions (Guo et al., 2015; de Wit et al., 2015; Sanborn et al., 2015), highlighting the importance of CTCF orientation in chromatin architecture, although the prevailing absence of new interactions between CTCF pairs now brought into convergent orientation, and the existence of CTCF-independent TADs (Nora et al., 2017; Taylor et al., 2022) suggests that other factors participate in TAD formation (Sikorska and Sexton, 2020).

Another feature predominantly enriched at TAD boundaries is active genes (Dixon et al., 2012; Sexton et al., 2012; Bonev et al., 2017). *A priori*, RNA polymerase promoter binding and subsequent tracking along chromatin during transcription may cause topological disruption of the underlying DNA fiber (Lavelle, 2014). It has thus been proposed that RNA polymerase binding and/or transcription may organize TADs, either directly or by acting as a CTCF-independent roadblock to loop extrusion (Brandão et al., 2019; Banigan et al., 2022). However, conflicting reports suggest that it remains unclear to what extent transcription modulates chromatin architecture, and whether any such effects are direct. For example, the inactive X chromosome lacks TADs globally except around the few genes escaping silencing (Giorgetti et al., 2016), yet transcriptionally inert sperm chromosomes maintain essentially all TADs (Du et al., 2017; Jung et al., 2017). One possible explanation for the discrepancy is that the non-coding RNA *Xist* excises cohesin from the inactive X chromosome (Minajigi et al., 2015), so loop extrusion processes could be perturbed chromosome-wide independently of transcriptional effects. TADs arise in early embryogenesis of mammals and flies, coinciding with the onset of zygotic gene activation, but the process is largely unaffected on treatment with drugs inhibiting transcription (Du et al., 2017; Hug et al., 2017; Ke et al., 2017). A different study in a cell line reported that TADs were weakened on similar pharmacological transcription inhibition (Barutcu et al., 2019). Looking more mechanistically, a controlled study of cohesin-mediated loop extrusion showed that this ATP-dependent process can occur independently of ongoing transcription (Vian et al., 2017), but this does not preclude other means of architectural modulation. For example, transcription, among other mechanisms, appears to influence cohesin loading onto chromatin (Busslinger et al., 2017), so may indirectly influence loop extrusion processes in some contexts. Alternatively, viral-induced extension of transcription beyond usual termination sites was found to disrupt TADs, perhaps due to removal of CTCF by the engaged RNA polymerase and hence loss of TAD boundary function (Heinz et al., 2018). As ultra-high-resolution chromatin interaction (micro-C) maps of mammalian chromosomes became available, small TAD-like domains at the level of single expressed genes began to be discerned (Hsieh et al., 2020), reminiscent of what was previously observed in yeast (Hsieh et al., 2015) and transcription-linked “compartmental domains” described in Drosophila (Rowley et al., 2017). This suggests two, non-mutually exclusive means by which gene activity can modulate chromatin architecture. Firstly, since chromatin tends to compartmentalize into co-associated active “A” compartments, separate from co-associated silent “B” compartments (Lieberman-Aiden et al., 2009), a process which is independent or perhaps even antagonistic to loop extrusion-mediated TAD formation (Nora et al., 2017; Rao et al., 2017; Schwarzer et al., 2017), small active genes within large regions of inactive chromatin could indirectly form a domain boundary by disrupting its resident B compartment (Rowley et al., 2017). Secondly, RNA polymerase tracking could directly compact the underlying transcription unit to generate its own sub-TAD, reminiscent of previous reports that the 5’ and 3’ ends of active genes interact to assure transcription directionality (Tan-Wong et al., 2012). In support of both of these phenomena, analysis of micro-C data in mouse embryonic stem cells (ESCs) revealed that intra-gene contact frequency correlates with RNA polymerase occupancy of the gene, and that RNA polymerase binding, while being a poor predictor of TAD boundary location, is a reasonably good predictor of TAD boundary strength (Hsieh et al., 2020). Despite these promising findings, the most direct tests of transcription perturbation (not just those relying on inhibitor drugs, which may have secondary effects) have shown rather limited effects on chromatin structure. Acute ablation of any of the three RNA polymerases with an auxin-inducible degron had negligible effects on any feature of genome architecture, including TADs (Jiang et al., 2020). The resolution of this study may not have been high enough to discern subtle effects, and a similar experimental approach showed that whereas interphase chromatin was indeed largely unaltered, resetting of TADs just after mitosis was affected by loss of RNA polymerase II (Zhang et al., 2021). As an alternative means of testing the direct effect of transcription on chromatin topology, another study used CRISPR-activation (CRISPRa; Gilbert et al., 2013) to ectopically induce two specific genes in ESCs: *Sox4* and *Zfp608* (Bonev et al., 2017). These genes were found to become new TAD boundaries on differentiation to neural precursors, concomitant with their transcriptional activation, although their induction in ESCs was unable to cause any topological changes at the target loci. Collectively, this body of work suggests that, despite extensive correlation between TAD organization and gene activity, it remains unclear whether there is a direct causative role for transcription in chromatin topology, and any such links are likely to be subtle and limited to specific genomic contexts.

In this study, we assessed the developmental dynamics of TAD organization, using mouse thymocyte development as a model system. Although TAD structures were largely conserved, in line with previous lower-resolution studies, we observed some specific TAD remodeling events which coincided with transcriptional changes. Notably, we observed apparent tracking of a broadened TAD border at the *Bcl6* gene concomitant with extension of pause-released RNA polymerase into the gene body, and cases of sub-TADs corresponding to single activated genes, lending further support to the previously mentioned models by which transcription could modulate chromatin topology. Most importantly, for a tested gene, *Nfatc3*, CRISPRa-induction of the gene in ESCs was sufficient to recapitulate the thymocyte TAD remodeling event, providing direct evidence that transcription can be a driver of chromosome folding in certain contexts.

## Results

### High-resolution chromatin architecture at key thymocyte genes

To assess at higher resolution whether TADs may be selectively remodeled around genes that are differentially expressed on developmental transitions, we performed Capture Hi-C (Dryden et al., 2014; Franke et al., 2016) on mouse CD4^-^ CD8^-^ CD44^-^ CD25^+^ (double negative; DN3) and CD4^+^ CD8^+^ (double positive; DP) thymocytes, representing cells just initiating and cells just after the beta-checkpoint for productive T cell receptor beta-chain rearrangement (Carpenter and Bosselut, 2010). We used tiled capture oligonucleotides spanning *Dpn*II fragments across eight ∼600 kb regions, centered on key thymocyte genes located close to TAD borders, identified from high-resolution Hi-C maps generated in mouse embryonic stem cells (ESCs) (Bonev et al., 2017): three that are upregulated on DN3-to-DP transition (*Bcl6, Nfatc3, Rag1*), three that are downregulated (*Cdh1, Il17rb, Pla2g4a*), and two whose expression are unchanged (*Cd3, Zap70*) (**Fig S1A** and **Table S1**). We also performed conventional Hi-C on DN3 and DP cells for genome-wide but lower-resolution assessment of any chromatin topology changes (overview of datasets in this study given in **Table S2**). Capture Hi-C was additionally performed on ESCs to provide a reference point and comparison with high-resolution conventional Hi-C data (Bonev et al., 2017). As expected, the Capture Hi-C provided improved resolution at the target regions compared to previously generated Hi-C maps for thymocytes (Hu et al., 2018), even though the latter dataset had ∼4-fold greater sequencing depth (**Fig 1A**). Biological replicates were highly correlated (Spearman correlation coefficient ≥ 0.95; **Fig S1B-C** and **Table S3**) and were pooled for improved resolution in subsequent analyses. The resultant maps were of sufficient quality to resolve chromatin interactions between promoters and putative enhancers (at *Bcl6*), and between CTCF-bound sites (at *Rag1*), further confirmed by 4C-seq analysis (**Fig 1B** and **S1D**). Visual inspection of the Capture Hi-C maps showed that thymocytes seem to have better defined TADs and more heterogeneous structures than ESCs (**Fig 1B**), in line with observations that ESC chromatin is generally more open and “plastic” than differentiated cells (Gaspar-Maia et al., 2011). Observing the contact strength decay of the Capture Hi-C datasets with increasing genomic distance reinforces these trends by showing greater variability of contact strength at a given genomic separation within thymocytes (**Fig 1C**). Further, the cis-decay profiles for both Capture Hi-C and, at lower resolution, the genome-wide Hi-C, show that DP chromatin is globally more compact than the other cell types, in agreement with previous observations (Rawlings et al., 2011). Overall, these results validate the quality of the Capture Hi-C datasets for subsequent in-depth comparisons across cell types.

**Figure 1.**
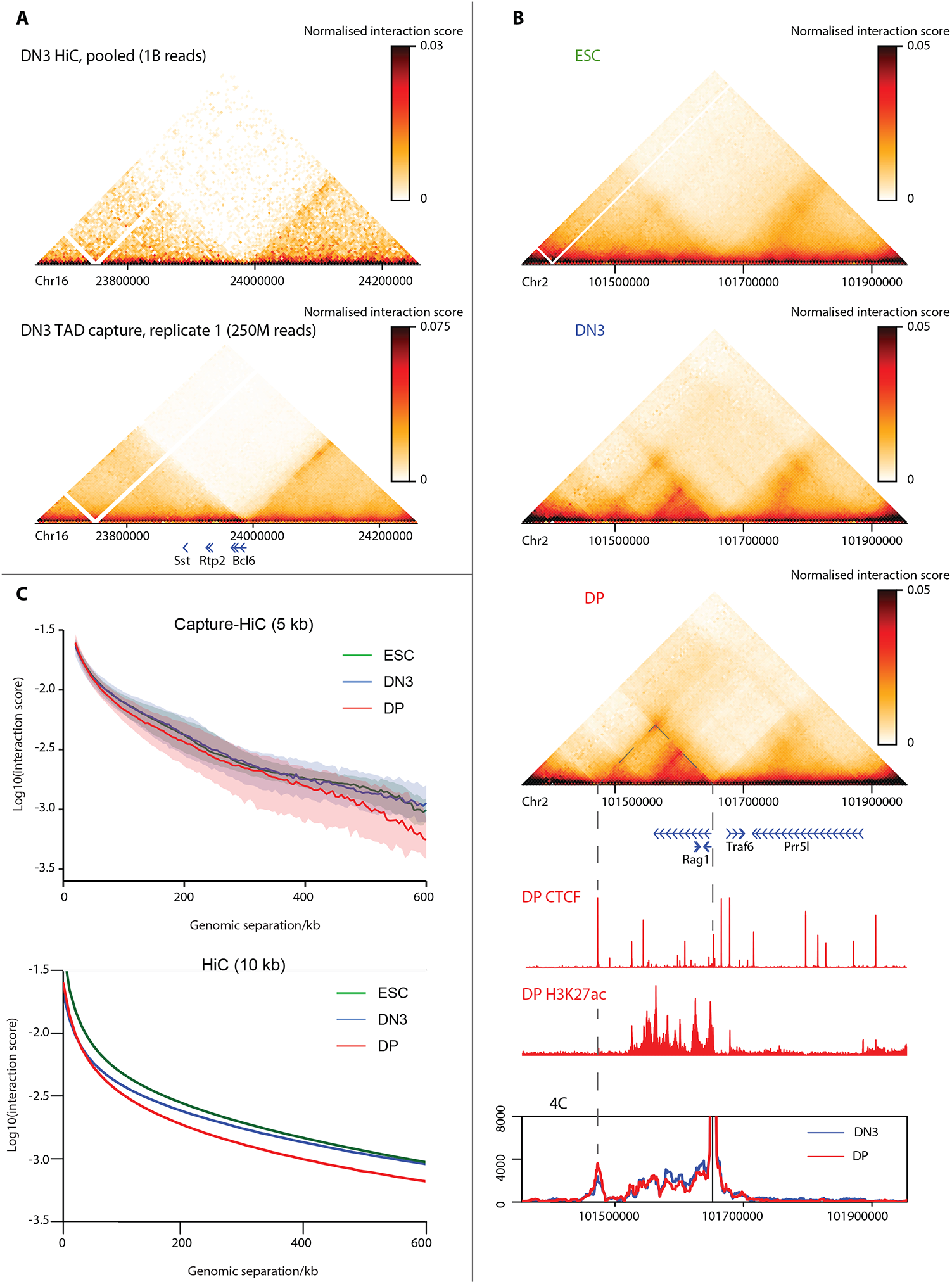
High-resolution interrogation of TADs in thymocytes with Capture Hi-C. A) Maps of DN3 chromatin interactions for a ∼600 kb region around the *Bcl6* gene from pooled Hi-C (from Hu et al., 2018; ∼1 billion reads; top) or one replicate of Capture Hi-C (∼250 million reads; bottom) are shown at 10 kb and 5 kb resolution, respectively. Positions of genes are denoted underneath the maps. B) Pooled Capture Hi-C interaction maps for a ∼600 kb region around the *Rag1* gene are shown for ESCs, DN3 and DP cells at 5 kb resolution. Below, in order, are shown positions of genes, and ChIP-seq tracks for CTCF and H3K27ac in DP cells. Bottom: 4C-seq profiles, using the *Rag1* promoter as bait, performed in DN3 (blue) and DP (red) cells. Dashed lines denote position of *Rag1* promoter and downstream CTCF site, which form an interaction in thymocytes, as shown on the Capture Hi-C and 4C maps. C) Plot of median normalized interaction score against genomic separation from the pooled Capture Hi-C (top) and Hi-C (bottom) datasets, at 5 kb and 10 kb resolution, respectively, for ESCs (green), DN3 (blue) and DP (red) cells. For the Capture Hi-C data, shading denotes the inter-quartile range to indicate the variability of these *cis*-decay distributions.

### TAD structure is largely conserved but with cell type-specific differences

Visual inspection of Capture Hi-C maps around the target regions suggested that TAD structure was largely conserved across the studied cell types, and particularly in between the two thymocyte populations, in line with lower-resolution studies (Dixon et al., 2015; Hu et al., 2018). To compare TAD organization more systematically, we computed insulation scores (Crane et al., 2015) at 5 kb resolution within the regions targeted by capture oligonucleotides. Compared to other methods which just call the positions of TADs (Forcato et al., 2017), insulation has the advantage of giving a TAD border “score” for all genomic intervals. Since TAD-like chromosomal domains are known to be nested and hierarchical (Fraser et al., 2015; Zhan et al., 2017), we computed insulation over multiple sliding window widths to alter sensitivity to domains of different size. In agreement with previous observations that TADs appear to be better defined in differentiated cells than in pluripotent cells, insulation scores are more homogeneous in ES cells, whereas more striking insulation score minima (representing strongly insulating TAD borders) and maxima (representing the centers of well-defined folded domains) are apparent in thymocytes (**Fig 2A**). Over a wide range of window sizes used to compute insulation score, the more homogeneous distribution in ES cells is significantly different to those of the thymocytes (two-sided Kolgorov-Smirnov test; **Table S4**). However, the genomic location of TAD border candidates is well conserved across cell types, since insulation scores are overall very well correlated (Spearman correlation coefficient between 0.71 and 0.91; **Fig 2B** and **Table S4**). These observations are supported by analysis of the lower-resolution Hi-C data, both at the specified capture regions and genome-wide (**Fig S2A-B** and **Table S5**). Thus on first analysis, TAD (border) *location* appears to be well conserved across cell types, but overall, TAD *strength* is higher in differentiated cells than in pluripotent ESCs.

**Figure 2.**
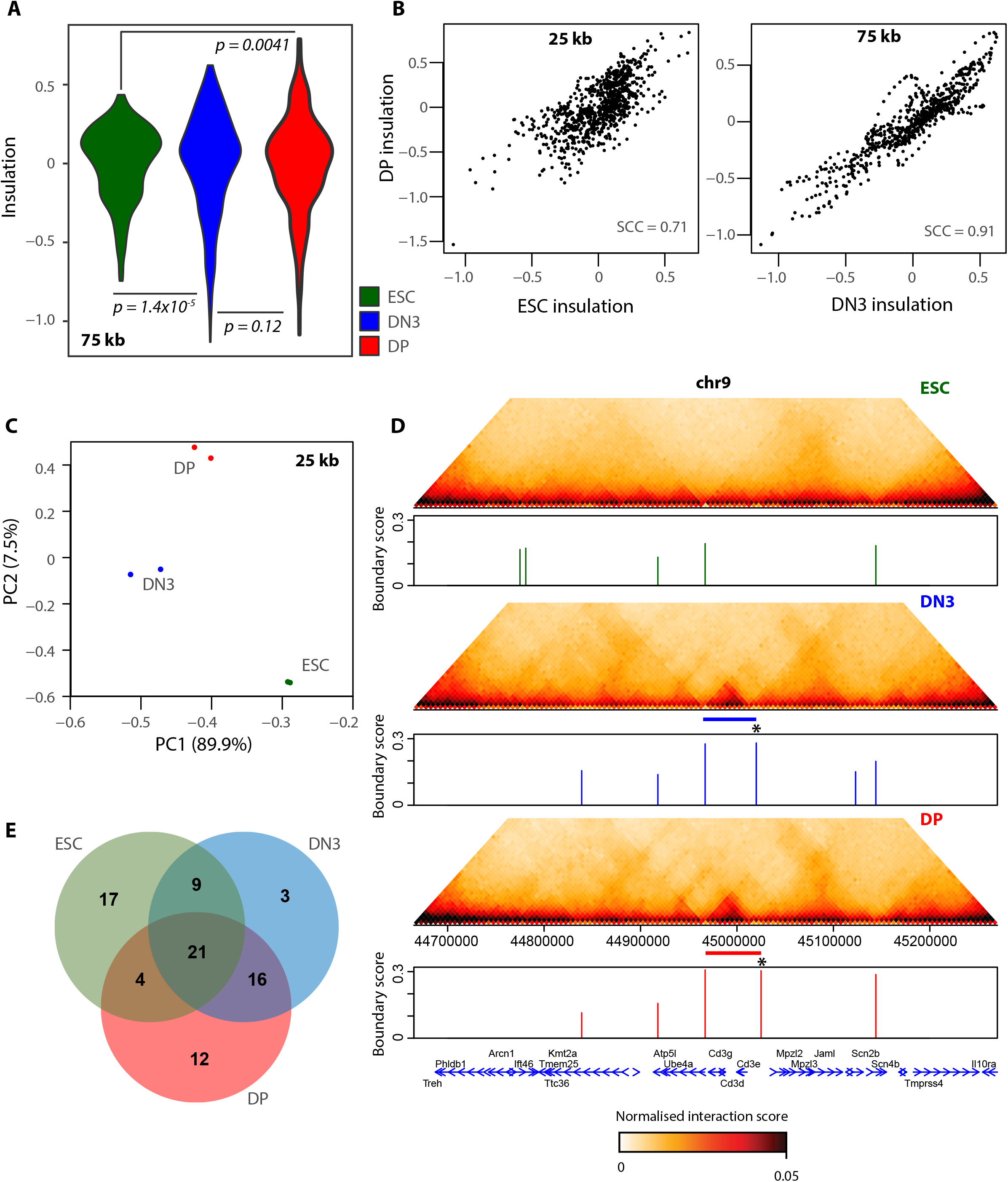
TADs are largely conserved, with cell type-specific differences. A) Violin plots for distributions of insulation scores computed on pooled Capture Hi-C datasets at 5 kb resolution, using a 75 kb (15 bin) window, on ESCs (green), DN3 (blue) and DP (red) cells. The more homogeneous ESC distribution is significantly different to DN3 (*p* = 1.4×10^−5^) and DP (*p* = 0.0041) cells, but the two thymocyte distributions are not significantly different to each other (*p* = 0.12; Kolgorov-Smirnov test). B) Scatter plots of insulation scores computed on pooled Capture Hi-C datasets at 5 kb resolution, using a 25 kb (5 bin; left) or 75 kb (15 bin; right) window, comparing DP cells with ESCs (left) or DN3 cells (right). Spearman correlation coefficients (SCC) are denoted. C) Plot of first two principal components for insulation scores computed on biological replicates of Capture Hi-C datasets at 5 kb resolution, using a 25 kb (5 bin) window for ESCs (green), DN3 (blue) and DP (red) cells. D) Pooled Capture Hi-C maps interaction maps for a ∼600 kb region around the *Cd3e* gene are shown, just above plots showing the positions and scores of called TAD boundaries in ESCs (green), DN3 (blue) and DP (red) cells. Positions of genes are shown below. Blue and red bars denote the position of a thymocyte-specific TAD, which is accompanied by the gain of a boundary at the 5’ end (denoted by asterisk). E) Venn diagram showing overlapping of called TAD boundaries for the pooled Capture Hi-C datasets of ESCs (green), DN3 (blue) and DP (red) cells.

To explore further whether there are any cell type-specific differences in TAD organization, we performed principal component analysis on the insulation scores. Across all tested insulation score window sizes, biological replicates clustered closely together, and the three cell types were clearly distinct from each other, suggesting that cell type-specific TAD patterns are indeed present (**Fig 2C** and **S2C)**. Such differences quantified by the principal component analysis could be due to gain/loss of specific borders to generate new TADs, altered strengths of pre-existing TADs (which has been observed for some genes at early thymocyte transitions; Hu et al., 2018), and/or accumulated small but reproducible changes dispersed across the studied genomic regions, which individually make no significant difference to the Hi-C maps. Since insulation scores from biological replicates clustered so closely together, we pooled the Capture Hi-C replicates and computed higher-resolution insulation scores over 2 kb genomic bins, finding strong local minima conserved across different sliding window sizes to call the more robust TAD borders. We identified ∼50 borders within the captured regions for each cell type (92 total; **Table S6**). In line with the above observations, many of the strongest borders were conserved throughout development, but cell type-specific TADs were apparent, in particular when comparing thymocytes to ESCs (**Fig 2D,E** and **Fig S2D**). As may be expected from their developmental proximity, the two thymocyte populations had more TAD borders in common (DN3-DP Jaccard index 0.57) than with ESCs (DN3-ESC Jaccard index 0.43; DP-ESC Jaccard index 0.32). Applying an analogous TAD border calling method to the Hi-C data at lower (10 kb) resolution gave similar results genome-wide (**Fig S3A-C**; **Table S7**), although the separation between thymocytes and ESCs was not as pronounced. This may be due to the focus of the Capture Hi-C around thymocyte-specific genes, which might be expected to have greater architectural differences than more general, housekeeping genes. Overall, we find a general conservation of TAD organization between different cell types, although higher-resolution analysis can uncover cell type-specific insulation patterns, sometimes associated with variant TAD borders.

### Pause-released polymerase may broaden TAD boundaries

We next further investigated the cell type-specific chromatin architectures uncovered by the Capture Hi-C approach. When comparing ESCs to thymocytes, we found cases of thymocyte-specific sub-TADs arising from establishment or strengthening of boundaries near the promoter of genes expressed specifically in thymocytes, both in Capture Hi-C (e.g. the *Cd3* cluster; **Fig 2D)** and lower-resolution Hi-C (e.g. *Runx1*; **Fig S3C**) maps. These new boundaries were accompanied by increased binding of CTCF near the promoter during thymocyte development (**Fig S3D**). For both genes, the downstream boundary appeared to be maintained across the studied cell types, corresponding with a conserved CTCF binding site. Based on the loop extrusion model, these developmental architectural changes between ESCs and thymocytes could be explained solely by differential CTCF binding. Whether underlying gene activation could be a cause or consequence of 3D genome structure, if functionally linked at all, thus remains unclear. However, interrogating differences between DN3 and DP chromatin structures, which have very similar CTCF binding profiles, uncovered potentially more direct links between transcription and chromatin architecture. Firstly, a strong, conserved TAD border at the *Bcl6* promoter becomes “leakier” in DP cells where the gene is highly expressed: an additional weak “stripe” extending the upstream TAD indicates the presence of a secondary broader border, adjacent to the major sharp boundary at the promoter (see rectangle in **Fig 3A**; **Fig S4A**,**B**). Notably, the TAD “extension” corresponds with the *Bcl6* coding region, and ChIP-seq data show that RNA polymerase II is paused at the promoter/major TAD boundary in DN3 cells while being released to fully transcribe *Bcl6* in DP cells. These Capture Hi-C data are thus consistent with a model whereby bound and/or engaging RNA polymerase can act as a topological boundary itself, perhaps by stalling loop extrusion processes, as has been reported in bacteria (Brandão et al., 2019) and proposed in a parallel study for mammalian cells (Banigan et al., 2022). In DN3 cells, the accumulation of RNA polymerase at the paused promoter could define a clear boundary, whereas the exact position of the elongating polymerase varies within each cell of a fixed DP population, generating a broader, more blurred boundary in the average map. While an attractive model, RNA polymerase II is absent from the inactive *Bcl6* gene in ESCs, where a relatively strong but somewhat broader TAD border is nevertheless maintained at the promoter, so other mechanisms must also contribute to architectural maintenance at this locus (**Fig 3B**). Conserved CTCF binding at the promoter would be expected to be such a mechanism, although homozygous deletion of the major binding site had mild effects on locus architecture (**Fig S5**), in line with other studies suggesting that TADs are built up cooperatively from multiple elements (Despang et al., 2019; Rodríguez-Carballo et al., 2017; Taylor et al., 2022).

**Figure 3.**
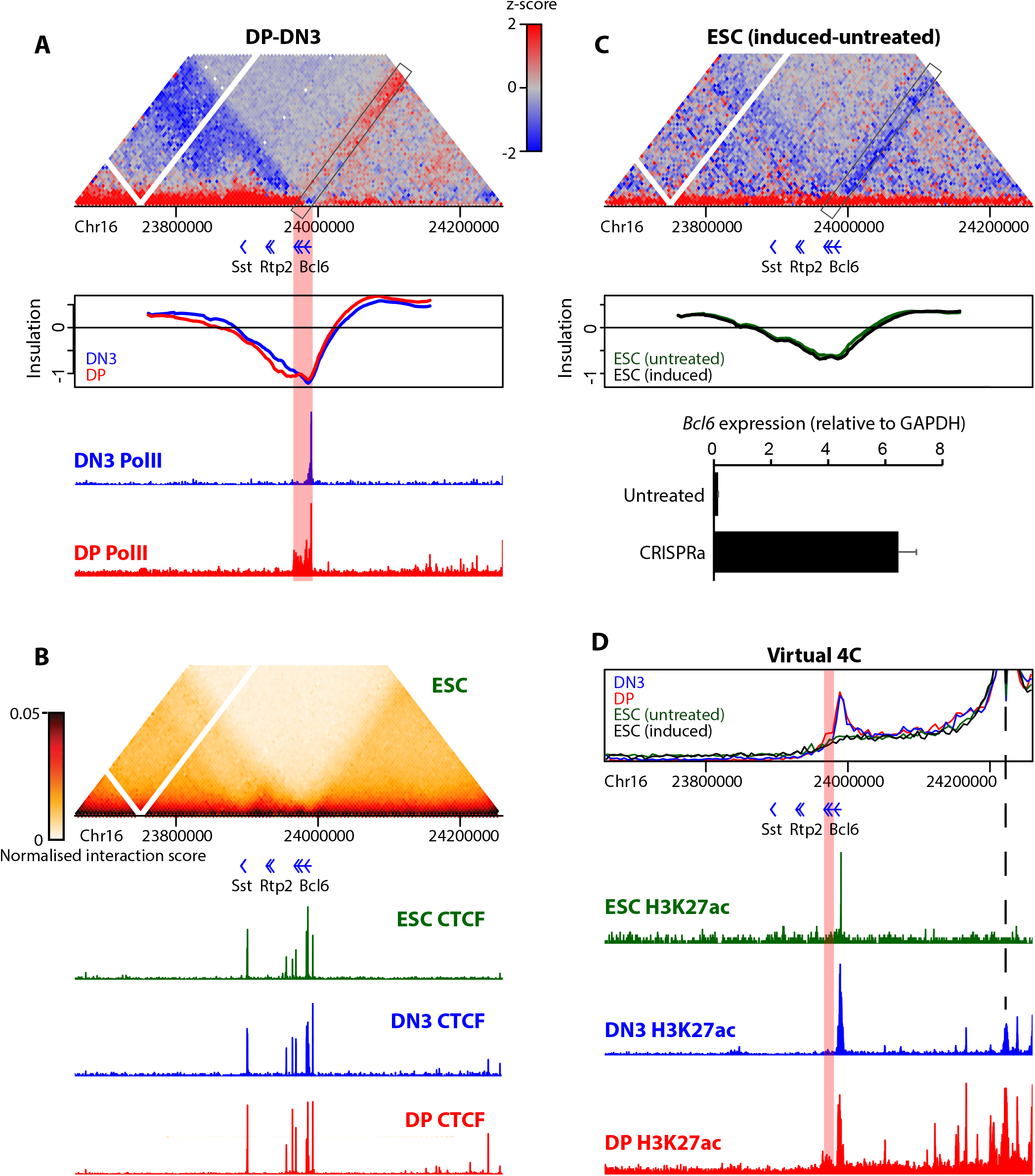
Broadening of TAD border at Bcl6 gene in DP cells tracks with RNA polymerase pause release. A) Differential Capture Hi-C map for a ∼600 kb region around the *Bcl6* gene is shown at 5 kb resolution, comparing interactions between DN3 and DP cells. Shown below, in order, are the positions of genes, the insulation scores in DN3 (blue) and DP (red) cells, computed on pooled datasets at 2 kb resolution with an 80 kb (40 bin) window, and ChIP-seq tracks for RNA polymerase II in DN3 (blue) and DP (red) cells. The rectangle on the differential map highlights the “stripe” of increased interactions for DP cells encroaching into the *Bcl6* gene body. The red rectangle shows that this region corresponds to persistent low insulation scores beyond the local minimum and extended RNA polymerase II engagement at the gene body in DP cells. B) Pooled Capture Hi-C interaction map for the same region as A) is shown for ESCs at 5 kb resolution, above ChIP-seq tracks for CTCF in ESCs (green), DN3 (blue) and DP (red) cells. C) Differential Capture Hi-C map for the same region as A), comparing interactions for untreated ESCs and those where *Bcl6* is induced by CRISPRa. Shown below, in order are the insulation scores in untreated (green) and CRISPRa-treated (black) ESCs, computed on pooled datasets at 2 kb resolution with an 80 kb (40 bin) window, and bar chart showing normalized *Bcl6* expression levels in untreated and CRISPRa-treated cells determined by qRT-PCR (two biological replicates). D) Virtual 4C plots derived from pooled Capture Hi-C datasets, using an upstream putative DP enhancer/H3K27ac peak as bait (position given by dashed line), for DN3 (blue), DP (red), untreated ESCs (green) and CRISPRa-treated ESCs (black). Shown below are the positions of genes and ChIP-seq tracks for H3K27ac in ESCs (green), DN3 (blue) and DP (red) cells.

To directly test the effect of transcription on TAD border extension, we performed CRISPRa (Gilbert et al., 2013) to recruit the activator VP64-p65-Rta (VPR; Chavez et al., 2015) specifically to the *Bcl6* promoter in ESCs. Despite a greater than 50-fold induction of the gene, there were negligible effects on the insulation score profile, and actually a slightly reduced interaction of the *Bcl6* gene body with upstream regions (**Fig 3C** and **Fig S4C**,**D**). Transcription of the gene therefore does not appear to directly affect TAD insulation at this gene. Alternatively, distal enhancer interactions may also play a role in TAD definition (Taylor et al., 2022), and they have previously been reported to track from the promoter into the gene body, presumably accompanying the progress of the engaged RNA polymerase (Lee et al., 2015). In line with this, virtual 4C from a putative interacting enhancer located ∼230 kb upstream of *Bcl6* (denoted by a peak of acetylation on histone H3 at lysine-27; H3K27ac) shows thymocyte-specific interactions with the *Bcl6* promoter, with DP-specific persistence of interaction into the downstream coding region (**Fig 3D**). The enhancer is inactive and non-interacting in ESCs, so does not affect TAD organization regardless of *Bcl6* expression. It is worth noting that DN3 and DP cells have an identical positioning of the local insulation score minimum, at the *Bcl6* promoter, with an equivalent score in both thymocyte populations. Instead, the insulation score stays consistently lower in the *Bcl6* coding sequence in DP cells (**Fig 3A**). Such subtle insulation changes are thus missed from conventional TAD boundary calling approaches, so it is unclear whether the TAD border broadening we observed at the *Bcl6* locus is a common occurrence in the genome. We did not observe this in any other region interrogated by the Capture Hi-C, and the resolution of our Hi-C data was insufficient to detect such subtle changes at the *Bcl6* gene or other loci in thymocytes.

### Transcription can directly remodel TADs

An additional type of topological change we observed when comparing thymocytes was the establishment of sub-domains encompassing single genes, specifically in the cell type where the gene is highly expressed. In the regions targeted by Capture Hi-C, we observed a DP-specific sub-TAD at the *Nfatc3* gene (**Fig 4A** and **Fig S6A**) and a DN3-specific sub-TAD at the *Tmem131* gene (**Fig 4B** and **Fig S6B**), neither of which are accompanied by altered CTCF recruitment. To test whether transcription can be a direct driver of such chromatin architecture, we used CRISPRa to induce *Nfatc3* ∼4-fold in ESCs where the sub-TAD is absent (**Fig 4C** and **Fig S6C**). Despite absolute expression of *Nfatc3* being ∼3-fold weaker than what was induced at *Bcl6*, a topological domain encompassing the *Nfatc3* gene was identified in the Capture Hi-C map, providing to our knowledge the first direct evidence that transcription could be instructive in TAD formation. Notably, the borders of the *Nfatc3* sub-TAD are also local minima of insulation score in conditions where the gene is silent, and either developmental (DN3-to-DP transition) or ectopic (CRISPRa in ESCs) induction does not change insulation score at these “proto-boundaries”. Instead, the overall intra-domain insulation score is increased, suggesting that transcriptional induction does not impede chromatin interactions between domains *per se*, but rather reinforces intra-domain contacts or compaction. For *Tmem131*, the insulation scores of the boundaries did differ somewhat between thymocyte stages, but the increase in intra-TAD contacts was much more pronounced. These results are reminiscent of previous observations of triptolide-sensitive single-gene domains in Drosophila (Rowley et al., 2017) and of a correlation between RNA polymerase occupancy and intra-gene contacts in ESCs (Hsieh et al., 2020) in ultra-high-resolution Hi-C or Micro-C maps, although neither study had assessed whether domains were directly formed as a consequence of transcription. Analysis of the thymocyte Hi-C datasets, restricted to sufficiently long genes to accommodate for the resolution limit, also showed an overall positive correlation between RNA polymerase occupancy and intra-gene contact strength (**Fig 4D**), quantitatively very similar to what was observed in ESCs (Spearman correlation coefficient of 0.60 (Hsieh et al., 2020), compared to DN3 (0.63) and DP (0.57) in this study). When clustering genes into four groups based on transcriptional output, metagene analysis confirms the gradual increase of intra-gene compaction by augmenting gene expression (**Fig 4E**). Additionally, the most highly expressed thymocyte genes also demonstrated stronger interactions between transcription start sites and transcription termination sites in metagene analysis (rightmost column in **Fig 4E**), in line with previous suggestions of active gene looping events (Tan-Wong et al., 2012), although the resolution is insufficient to distinguish between point-to-point interactions between gene termini and general compaction of the entire gene body to a more homogeneous domain. These and the previously reported correlations apply to comparisons of gene sets within the same cell type. An advantage of our experimental setup is that we can also compare architectures for the same genes in different, but developmentally very close, cell types. When comparing expression and intra-gene compaction *changes* between DN3 and DP cells, we also find a significant but much weaker positive correlation (Spearman correlation coefficient 0.24) (**Fig 4F**). This suggests that, globally, gene upregulation can remodel local chromatin domains, but is not likely to be a universal occurrence. Such a context-dependent architectural remodeling is supported by previous CRISPRa experiments which did not alter local chromatin topology (Bonev et al., 2017), and a counter-example in our own Capture Hi-C data of the *Il17rb* gene, which appears to fit the same criteria of *Tmem131* to form a DN3-specific domain (sufficiently long gene forming a cell type-specific RNA polymerase-bound domain flanked by local insulation score minima) but demonstrates no obvious differences in chromatin interactions (**Fig S6D**). Trials of multiple gRNAs failed to induce *Tmem131* in ESCs in CRISPRa trials, so we were unable to assess whether this domain was also directly remodeled by transcription. Overall, our Capture Hi-C and CRISPRa results at *Nfatc3* strongly support that direct TAD remodeling by transcription can occur, but with an apparent context-dependence which is hidden from previous global analyses.

**Figure 4.**
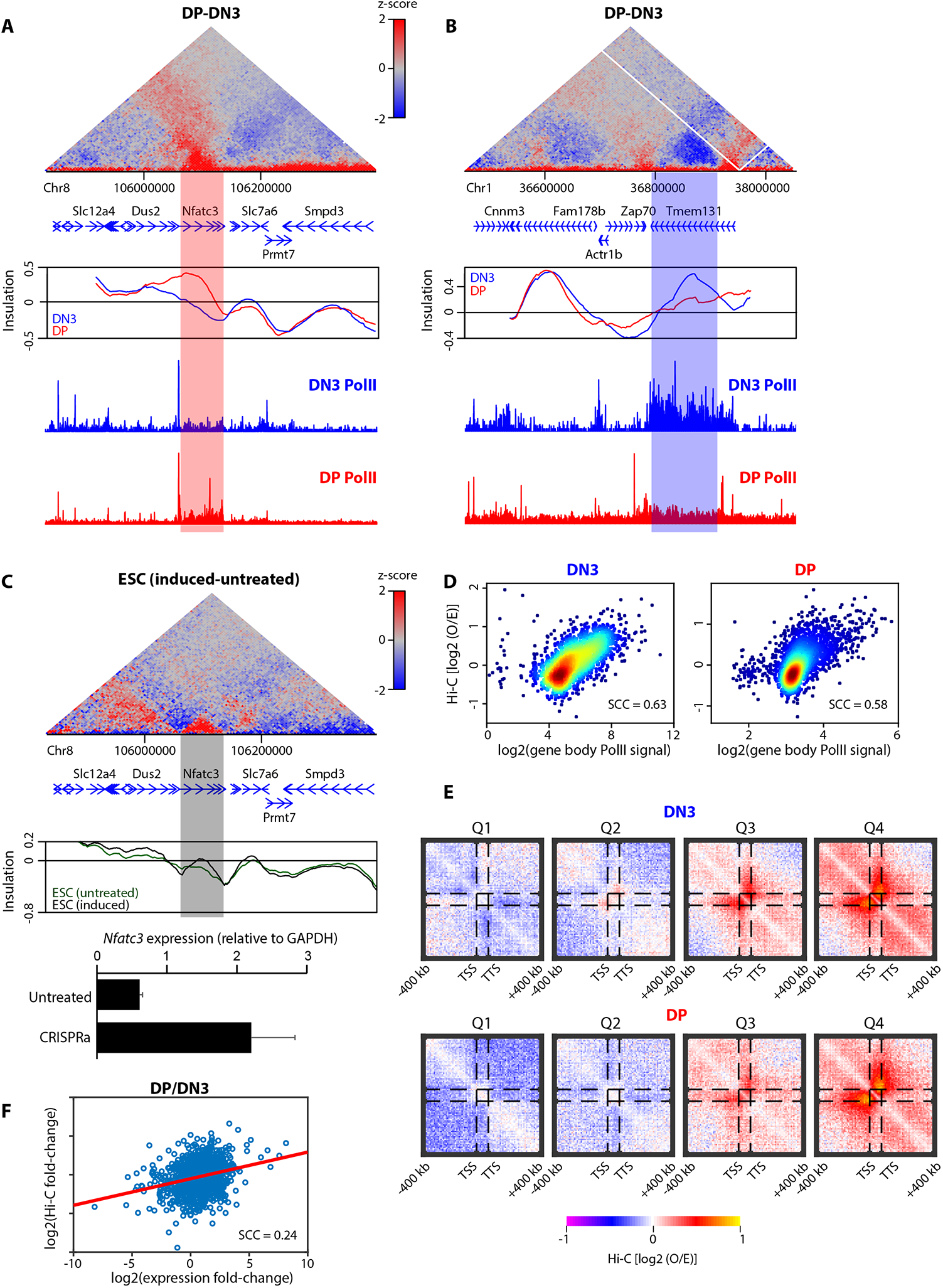
Transcription can directly remodel TADs around activated genes. A) Differential Capture Hi-C map for a ∼500 kb region around the *Nfatc3* gene is shown at 5 kb resolution, comparing interactions between DN3 and DP cells. Shown below, in order, are the positions of genes, the insulation scores in DN3 (blue) and DP (red) cells, computed on pooled datasets at 2 kb resolution with a 50 kb (25 bin) window, and ChIP-seq tracks for RNA polymerase II in DN3 (blue) and DP (red) cells. The red rectangle shows the DP-specific domain which corresponds to altered intra-domain insulation profile and increased RNA polymerase II engagement. B) Differential Capture Hi-C map for a ∼600 kb region around the *Tmem131* gene is shown at 5 kb resolution, comparing interactions between DN3 and DP cells. Shown below, in order, are the positions of genes, the insulation scores in DN3 (blue) and DP (red) cells, computed on pooled datasets at 2 kb resolution with an 80 kb (40 bin) window, and ChIP-seq tracks for RNA polymerase II in DN3 (blue) and DP (red) cells. The blue rectangle shows the DN3-specific domain which corresponds to altered intra-domain insulation profile and increased RNA polymerase II engagement. C) Differential Capture Hi-C map for a ∼500 kb region around the *Nfatc3* gene the same region as A) is shown at 5 kb resolution, comparing interactions between untreated ESCs and those where *Nfatc3* is induced by CRISPRa. Shown below, in order, are the positions of genes, the insulation scores in untreated (green) and CRISPRa (black) cells, computed on pooled datasets at 2 kb resolution with a 50 kb (25 bin) window, and bar chart showing normalized *Nfatc3* expression levels in untreated and CRISPRa-treated cells determined by qRT-PCR (two biological replicates). The gray rectangle shows the induction-specific domain which corresponds to altered intra-domain insulation profile. D) Scatter plots for intra-gene contact score relative to RNA polymerase II occupancy in DN3 (left) and DP (right) cells. SCC is denoted. E) Pileup Hi-C maps at 10 kb resolution showing the cumulative interaction between transcription start sites and termination sites in DN3 (top) and DP (bottom) cells with genes stratified according to expression levels, from the lowest quartile (Q1; left) to the highest quartile (Q4; right). F) Scatter plot for the ratio of intra-gene contact score relative to the ratio of expression for genes between DN3 and DP cells. SCC is denoted.

## Discussion

In this work, we assessed whether TAD remodeling events accompanied mouse thymocyte maturation, focusing at higher resolution on key, differentially expressed genes located close to strong TAD boundaries, where any such structural changes were most likely to occur. In line with lower-resolution studies (Hu et al., 2018), the majority of TADs appeared unchanged, despite significant transcriptional changes in hundreds of genes. Conventional measures of TAD architecture, such as the insulation score, were highly similar, both between thymocytes and unrelated cell types such as pluripotent cells, suggesting that most chromatin architecture at this scale is somehow “hard-wired” (Dixon et al., 2015), fitting with evolutionary conservation of TAD positions at syntenic regions across species (Vietri Rudan et al., 2015). Physical models propose that TAD homeostasis may be largely explained by cohesin-mediated loop extrusion interfering with higher-order configurations such as co-association of active compartments (Nuebler et al., 2018). The latter are dependent on underlying gene activity and epigenetic state so are developmentally plastic, as observed (Dixon et al., 2015; Bonev et al., 2017), whereas loop extrusion barriers could be genetically coded, such as by CTCF binding sites, so in this model TADs would not be expected to change significantly across cell type. However, CTCF occupancy is not identical in all cell types and in any case does not account for all TADs (Nora et al., 2017; Taylor et al., 2022). Recent studies have shown that non-encoded features as diverse as MCM complex binding (minichromosome maintenance; most famous for replication origin licensing) and DNA double-strand breaks can also be barriers to loop extrusion (Arnould et al., 2021; Dequeker et al., 2022). Thus small modifications to the basic “loop extrusion versus compartmentalization” model could just as easily explain tissue-specific TAD structures. An important challenge in the field will be to identify more comprehensively the features, both those genetically encoded/invariant and those more variable and linked to genomic functions, which can obstruct loop extrusion, and to assess their relative contribution to steady-state chromatin architecture.

Higher-resolution studies (Bonev et al., 2017; Rowley et al., 2017; Hsieh et al., 2020), including this one, do identify tissue-specific variations in TADs; they were presumably overlooked in previous works because they are too subtle and/or at too small a scale to be resolved. It is important to note that the most commonly utilized method to find TAD boundaries, identifying insulation score minima, does not detect the most important changes that we observed in this study. Expression-linked sub-TADs on genes and cell type-specific broadened TAD borders maintained the same local insulation minima in the cell types tested; instead, the insulation maxima were increased, or lower insulation scores extended beyond the minima, respectively. There is thus scope for the development and wider application of alternative Hi-C analytical methods to find otherwise overlooked architectural changes, such as the recently described CHESS (Galan et al., 2020). The major feature correlating with TAD remodeling events in ours and others’ studies is gene activity, further adding to the conflicting evidence as to whether or not transcription can drive genome topology (see Introduction). We provide a very convincing case that transcription can indeed play an instructive role, by recapitulating the sub-TAD around the *Nfatc3* gene by transcriptional induction in ESCs. While compelling, this was found for only one gene, with other genes providing counter-examples in previous work (Bonev et al., 2017). Further counter-examples are also present in this study. First an excellent candidate gene, *Il17rb*, did not show any architectural differences between thymocyte populations. Second, *Bcl6* induction in ESCs caused no structural changes, although thymocytes did not demonstrate sub-TAD formation either, but instead a different mode of TAD remodeling. Our study thus highlights two major questions for the field. Firstly, how exactly does transcription affect TADs? The main, non-mutually-exclusive hypotheses already proposed are that bound RNA polymerase is a direct obstacle to loop extrusion (Brandão et al., 2019), and that transcribed units form small A compartments disrupting the B compartments within which they reside (Rowley et al., 2017). Our results do not exclude either model, and actually suggest that one may be prominent over the other in a context-dependent manner. In the case of *Bcl6*, conventional CTCF-mediated architecture appears to play a role, particularly in ESCs where there is no transcription. However, despite a similar CTCF binding profile at this locus for all three cell types studied, the boundary is much sharper in DN3 cells, corresponding exactly with the position of paused RNA polymerase. Release of this pause on full transcription in DP cells re-widens the TAD boundary again, which could be explained if the polymerase acted as a mobile loop extrusion barrier. Similar gene body-broadened boundaries have recently been reported on metagene analysis in a parallel study (Banigan et al., 2022), and we have recently proposed in fission yeast that paused or backtracking RNA polymerase may have different effects on condensin-mediated loop extrusion to elongating polymerase (Rivosecchi et al., 2021). Ectopic induction of the gene had no effect on local chromatin topology in ESCs, but the boundary was already rather wide, so the model is not necessarily disproved. However, the intriguing finding that the secondary boundary also tracks with apparent distal enhancer interactions raises the possibility of alternative gene looping mechanisms being involved. Further work on more loci at high resolution is required to tease out these different possibilities. On the other hand, transcriptional activation at the genes *Nfatc3* and *Tmem131* caused no apparent changes to the local insulation minima flanking these genes, which are much weaker candidate TAD boundaries than for *Bcl6*. Instead, the entire gene body appeared to form its own sub-domain, in line with previous reports in yeast and ESCs (Hsieh et al., 2015; Hsieh et al., 2020). In this case, there is no obvious means by which loop extrusion is being affected, since contacts between the gene and flanking regions do not appear to be changed. Rather, increased intragenic contacts suggest some looping or compaction of the gene unit. Since we were unable to fairly compare large-scale gene interactions with the rest of the chromosome in the Capture Hi-C experiments, it is unclear whether this local structural change is accompanied by a change in compartment. If it did, this would be consistent with the “compartmentalized domain” model proposed in Drosophila (Rowley et al., 2017), since most fly genes would be too short to easily observe any accompanying intragenic compaction. To see how these two models of polymerase-mediated architectural remodeling interplay, it will first be necessary to demonstrate in single-molecule biophysical studies whether or not the polymerase is able to obstruct loop extrusion, a formidable challenge.

The second major question arising is why are transcription-mediated TAD remodeling changes not more universally observed? Seeing as most of the transcription-mediated effects appear to be subtle, it is possible that they are masked at most loci by more dominant factors, such as CTCF-mediated TAD boundary definition. As more high-resolution micro- and Hi-C maps become available, especially in comparative studies of more developmentally related cell types, such as in this study, more cases of transcription-mediated changes may be discerned. In this case, we will be in a better position to assess which principles may influence TAD positioning and strength at each locus. In combination with more sophisticated single-molecule (Davidson et al., 2019) and live-imaging (Gabriele et al., 2022), going beyond basic principles to understanding the context-specific nature of chromatin architecture at specific loci is becoming a more achievable goal.

## Acknowledgments

We thank Susan Chan for sharing reagents and Alexandra Pekowska for sharing R code for Hi-C matrix balancing. Sequencing was performed by the IGBMC GenomEast platform, a member of the France Génomique consortium (ANR-10-INBS-0009). This study was made possible because of the IGBMC flow cytometry and molecular biology platforms. We thank PSMN (Pôle Scientifique de Modélisation Numérique) of the ENS de Lyon for computing resources. Work in the Sexton lab was supported by funds from the European Research Council (ERC) under the European Union’s Horizon 2020 research and innovation program (Starting Grant 678624 - CHROMTOPOLOGY), the ATIP-Avenir program, and the grant ANR-10-LABX-0030-INRT, a French State fund managed by the Agence Nationale de la Recherche under the frame program Investissements d’Avenir ANR-10-IDEX-0002-02. YBZ was additionally supported by la Region Grand Est. Work in the Jost lab was supported by funding from Agence Nationale de la Recherche (ANR-18-CE12-0006-03; ANR-18-CE45-0022-01).

## Author contributions

SC, YBZ and TS designed the study. SC, NK and CE performed Hi-C and Capture Hi-C, and AMM and NK performed 4C-seq. AMM, DK, MM and AM optimized and performed the CRISPRa experiments. YBZ and TS performed the Capture Hi-C analysis. HS and DJ performed metagene analysis on the Hi-C data. All authors contributed to the writing and critical reading of the manuscript.

## Methods

### Thymocyte isolation

Thymocytes were obtained from 4-week-old mouse thymus by FACS. Thymuses were dissected, gently homogenized and filtered through a 70 μm strainer in cold PBE (0.5% BSA and 2 mM EDTA in 1X PBS). To purify DP cells, ∼600 million cells were incubated with anti-mouse CD4-PE, anti-mouse CD8a-FITC and anti-mouse CD3e-APC antibodies (eBioScience), before washing with cold PBE, adding DAPI and purifying viable (DAPI^-^) PE^+^ FITC^+^ APC^+^ cells by FACS. To purify DN3 cells, ∼4 billion cells were first depleted of DP cells by incubating with rat anti-CD4 and rat anti-CD8 anti-sera (kind gift of Susan Chan), washing with cold PBE and binding the DP cells to Dynabeads-sheep anti-rat IgG (Invitrogen). The supernatant was centrifuged to harvest the cells, which were then incubated with anti-mouse CD4-FITC, anti-mouse CD8a-FITC, anti-mouse CD3e-FITC, anti-mouse B220-FITC, anti-mouse CD11b-FITC, anti-mouse Ly-6G(Gr-1)-FITC, anti-mouse NK1.1-FITC, anti-mouse CD44-APC and anti-mouse CD25-PE antibodies (eBioScience) and washed with PBE. DAPI was added and viable PE^+^APC^-^FITC^-^cells were purified by FACS.

### ESC culture

J1 mouse ESCs were grown on gamma-irradiated mouse embryonic fibroblast cells under standard conditions (4.5 g/L glucose-DMEN, 15% FCS, 0.1 mM non-essential amino acids, 0.1 mM beta-mercaptoethanol, 1 mM glutamine, 500 U/mL LIF, gentamicin), then passaged onto feeder-free 0.2% gelatin-coated plates for at least two passages to remove feeder cells before treatments.

### CRISPRa

Twenty million cell batches of J1 ESCs were transfected each with 20 μg dCas9-VPR vector (Addgene #63798) and 20 μg plasmid constructed by the IGBMC molecular biology platform (available on request), containing optimized scaffold for 4 gRNAs alongside a puromycin resistance gene and mCherry reporter (gRNA sequences are given in **Table S8**), with Lipofectamine 2000 according to the manufacturer’s instructions. After 24 h, transfected cells were treated with 3 μg/mL puromycin and 1 mg/mL G418, and after another 24 h, mCherry-positive cells were sorted by FACS before immediate processing.

### Deletion of major Bcl6 CTCF site

One million J1 ESCs were transfected with 1 μg of a plasmid constructed by the IGBMC molecular biology platform (available on request), containing optimized scaffold for 2 gRNAs alongside a Cas9-HF-EGFP construct and a puromycin resistance gene, with Lipofectamine 2000 according to the manufacturer’s instructions. After 3 days, transfected cells were treated with 3 μg/mL puromycin for 24 h, then with 1 μg/mL puromycin for a further 24 h, before sorting individual GFP-positive cells into 96-well plates, pre-prepared with feeder cells, with FACS. Colonies were amplified and clones with homozygous deletions were identified by genomic DNA extraction and PCR screening with different primer pairs flanking and inside the expected deletion region. Sequences of the deletion were confirmed by TA-cloning and sequencing the PCR products.

### RNA isolation and gene expression analysis by RT-qPCR

Total RNA was extracted using the Nucleospin RNA kit (Macherey Nagel) and cDNA was prepared using SuperScript IV (Invitrogen) with random hexamers according to the manufacturer’s instructions. Quantitative PCR was performed in a LightCycler 480 (Roche) using SYBR Green I master mix according to the manufacturer’s instructions. Expression was normalized to *Gapdh* (primer sequences are given in **Table S9**).

### Capture oligonucleotide design

The mouse genome (mm10) was digested *in silico* with *Dpn*II and fragments were filtered to include those ≥ 140 bp and with GC-content between 20 and 80%. To assess target mappability, the genome was split *in silico* into 50 bp fragments and re-mapped using Bowtie (Langmead et al., 2009) with a filter to only include uniquely mapping fragments. The *Dpn*II fragments were further filtered to only include those where ≥ 80% of the sequence is covered by this uniquely mapping reference point. Fragments within 600 kb of the target genes (**Table S1**) were retained and the 120 nt directly adjacent to the *Dpn*II sites were used for capture oligonucleotides (full capture design is given in **Table S10**), synthesized as a SureSelect library (Agilent).

### Capture Hi-C

Capture Hi-C was essentially performed as in (Karasu and Sexton, 2021). Five million aliquots of cells were fixed in 2% formaldehyde for 10 min, quenched with 125 mM cold glycine, then collected by centrifugation and washed with PBS. Cells were lysed in lysis buffer (10 mM Tris-HCl pH 8, 100 mM NaCl, 0.2% NP-40, protease inhibitor cocktail) on ice for 30 min and nuclei collected by centrifugation and resuspended in *Dpn*II restriction buffer (NEBuffer 2 for *Hind*III Hi-C). ESCs were permeabilized by treatment with 0.4% SDS for 1 h at 37°C, followed by 1.6% Triton X-100 for 1 h at 37°C. Thymocytes were permeabilized by treatment with 0.8% SDS for 20 min at 65°C, followed by 40 min at 37°C, then with 3.3% Triton X-100 for 1 h at 37°C. Nuclei aliquots were digested overnight at 37°C with 1500 U *Dpn*II (2000 U *Hind*III for *Hind*III Hi-C), then cohesive ends were filled in with biotin tags by incubating for 90 min at 37°C with 15 μM dATP, 15 μM dTTP, 15 μM dGTP, 15 μM biotin-14-dCTP and 25 U DNA polymerase I Klenow fragment. Nuclei were collected by centrifugation and *in situ* ligation was performed overnight at 16°C in 500 μL volumes of 1X ligase buffer (NEB) with 20,000 U T4 DNA ligase, before de-crosslinking overnight at 65°C with 0.75 mg/mL proteinase K. DNA was purified by RNase A treatment, phenol/chloroform extraction and isopropanol precipitation. Five microgram aliquots of DNA were sonicated to a fragment size of ∼100-400 bp in a Covaris sonicator E220 and bound to Steptavidin MyOne T1 beads, following the manufacturer’s instructions. The DNA was then end-repaired by incubating for 1 h at 20°C with 0.8 mM dNTPs, 0.3 U/μL T4 DNA polymerase, 1 U/μL T4 polynucleotide kinase and 0.1 U/μL DNA polymerase I Klenow fragment in ligase buffer, then A-tailed by incubating for 1 h at 37°C with 200 μM dATP and 0.2 U/μL Klenow fragment (3’-5’ exo^-^) in NEBuffer 2 (NEB), then Illumina PE adapter was added by incubating overnight at 20°C with 15 μM PE adapter and 2000 U T4 DNA ligase in ligase buffer, washing the beads with appropriate buffers in between each treatment. Hi-C material was then amplified from the beads with 7-9 cycles of PCR using Herculase II Fusion DNA polymerase (Agilent) according to the manufacturer’s instructions. Material at this stage was quantified on a BioAnalyzer (Agilent) and *Dpn*II and *Hind*III Hi-C material was sequenced (2×50 nt) on a HiSeq 4000 (Illumina) following the manufacturer’s instructions). For Capture Hi-C, 750 ng *Dpn*II Hi-C material was concentrated in a SpeedVac vacuum concentrator and then target capture was performed using the SureSelect XT Target Enrichment system (Agilent) with the custom-designed library (**Table S10**), following the manufacturer’s instructions and the library was then sequenced (2×50 nt) on a HiSeq 4000 (Illumina).

### 4C-seq

3C was performed on ESCs and thymocytes essentially as for the first steps of the Hi-C described above (up to DNA purification before sonication), except that the biotin fill-in step was omitted. The DNA was digested with 5 U/μg *Csp*6I at 37°C overnight, then re-purified by phenol/chloroform extraction and isopropanol precipitation. The DNA was then circularized by ligation with 200 U/μg T4 DNA ligase under dilute conditions (3 ng/μL DNA), and purified by phenol/chloroform extraction and isopropanol precipitation, before PCR amplification with primers containing Illumina adapter sequence (sequences given on **Table S11**). Excess primers were removed with SPRI beads, and the PCR products were assessed on a BioAnalyzer (Agilent) then sequenced (1×50 nt) with a HiSeq4000 (Illumina).

### Hi-C analysis

For convenience, Hi-C data were processed using the FAN-C suite of tools (Kruse et al., 2020), entailing read mapping to the mm10 genome assembly, mating pairs, attribution to restriction fragments with filtering of self-ligated fragments and PCR duplicates, matrix balancing, and computing insulation scores. For compatibility with Capture Hi-C plots, the data for the relevant regions were extracted using the *dump* tool, and visualized as for the Capture Hi-C matrices.

### Capture Hi-C preprocessing

To ensure that the captured regions were treated independently of the non-captured regions, the datasets were processed with custom scripts, as entailed in (Ben Zouari et al., 2019). The initial steps are essentially the same as FAN-C, entailing read mapping with Bowtie (Langmead et al., 2009) to the mm10 genome assembly, mating pairs, and attribution to restriction fragments with filtering of self-ligated fragments and PCR duplicates. Custom perl scripts filter the data to include only those where both ends of a paired read correspond to regions targeted by capture oligonucleotides, and to bin these sub-matrices to fixed genomic bins. Each captured sub-region was treated separately for matrix balancing with the Knight-Ruiz method (Rao et al., 2014; R code kindly shared by Alexandra Pekowska). All plots in this study visualizing (Capture)-HiC matrices alongside ChIP-seq tracks and insulation scores were generated by custom R scripts.

### 4C-seq analysis

4C-seq data were mapped to the mm10, processed, normalized and visualized using 4See (Ben Zouari et al., 2020), as in (Taylor et al., 2022).

### (Capture) Hi-C analysis

The cis-decay plot for **Fig 1C** was obtained from 10 kb Hi-C data with the *expected* function of FAN-C. For the 5 kb Capture Hi-C data, the cis-decay plot was derived directly from the balanced sub-matrices at the captured regions, obtaining the distribution of normalized interaction scores at different genomic separations, and plotting the median values ± the inter-quartile range.

Insulation scores were essentially computed as in (Crane et al., 2015). determined by counting interactions along sliding windows off the Hi-C diagonal, with lower counts/weaker interactions implying greater topological insulation and thus a greater likelihood of the existence of a TAD border. This has the additional advantage of giving a TAD border “score” for all genomic intervals, rather than simply calling TADs using a more complex set of (often arbitrary) parameters. The only tunable parameter required in computing insulation is the sliding window size, altering the sensitivity to TADs of different sizes within the folded hierarchy (Zhan et al., 2017). Local insulation score minima are candidate TAD boundaries, and their *boundary scores* are computed as the difference in insulation score between the minima at the boundary and the adjacent bins (the *delta* in Crane et al., 2015), with higher scores indicating sharper, stronger TAD borders. For 10 kb Hi-C data, insulation score was computed with the *insulation* tool of FAN-C, using sliding windows of 70, 100 and 150 kb (7, 10 and 15 bins). Local minima were subsequently identified with the FAN-C tool *boundaries*, using a window of 3 bins for computing the boundary score and with no initial threshold filter for minimal boundary score. For Capture Hi-C, insulation score was computed with custom R scripts on the balanced sub-matrices at 5 kb (for individual replicates) and 2 kb (for pooled data) resolutions, using sliding windows of 15, 25, 35, 50 and 75 kb (3, 5, 7, 10 and 15 bins), and 30, 40, 50, 80 and 100 kb (15, 20, 25, 40 and 50 bins), respectively. Windows of 3 bins were used for computing boundary scores for each of these insulation profiles. To identify the most robust TAD boundaries (e.g., as shown in **Fig 2D**), insulation score minima with a boundary score ≥ 0.1 in at least two different window sizes were maintained. Adjacent boundaries were merged and the overall boundary score for each boundary was calculated as the mean of the component scores which were ≥ 0.1. These final called borders were intersected using the *GenomicRanges* R package to derive the Venn diagrams (e.g. **Fig 2E**). Distributions of insulation score were plotted as violin plots (e.g. **Fig 2A**) using the *ggplot2* package of R, and were compared across cell type by two-tailed Kolgorov-Smirnov tests.

Biological replicates were compared by Spearman correlation coefficients of normalized interaction scores (for Capture Hi-C) and insulation scores (for Hi-C and Capture Hi-C). Principal component analysis was performed on the biological replicates of Capture Hi-C insulation scores using the *prcomp* function of R.

To derive the differential interaction maps (e.g. **Fig 3A**) the normalized scores of one matrix were subtracted from the other, and the difference was expressed as a z-score: (diff – mean(diff))/standard deviation(diff).

Virtual 4C plots were derived as one row from the relevant Capture Hi-C normalized interaction score submatrix, applying smoothing with a running mean over three bins.

### Metagene analysis

For all mouse genes of size ≥ 100 kb (≥10 bins in Hi-C), the intragenic interaction for each gene in each thymocyte population was computed as the median observed/expected score (defined as the Hi-C interaction normalized by the median Hi-C interaction level at the same genomic separation computed over the whole genome; see cis-decay curve in **Fig 1C**) for all intragenic pairwise bin-to-bin combinations. For the same gene set, the median RNA polymerase II ChIP signal within the gene body (taken over 10 kb bins) was also calculated and used to generate the scatter plots in **Fig 4D. Fig 4E** was made like in (Rowley et al., 2017) and (Hsieh et al., 2020). Genes ≥ 100 kb were grouped into four quartiles based on their expression levels, and, for each group, the median heatmap of observed/expected interaction scores are computed, by scaling genes to a pseudo-size of 10 bins to align all the transcription start and termination sites for the intra-gene part of heatmap, and by considering extensions of 400 kb (40 bins) beyond the genes (upstream and downstream) for the rest. In **Fig 4F**, for each gene, the log2 ratio of the intragenic interaction score in DP and DN3 is plotted versus the corresponding log2 ratio of gene expression levels (quantified by RNA-seq data).

### Epigenomic datasets

Publicly available thymocyte and ESC RNA-seq and ChIP-seq data were obtained from the Genome Expression Omnibus (see **Table S12**), and BigWig tracks were processed and visualized with the R package *rtracklayer*.

## Supplemental Information

**Supplemental Figure 1. High-resolution interrogation of TADs in thymocytes with Capture Hi-C**. A) ESC Hi-C maps (data taken from Bonev et al., 2017) are shown at 5 kb resolution for ∼600kb regions surrounding the genes *Rag1* (left) and *Pla2g4a* (right), showing a TAD border near these genes. Below are shown the positions of genes and RNA-seq tracks (non-strand-specific) from DN3 (blue) and DP (red) cells, showing differential expression of the target genes between the thymocyte populations. B) Capture Hi-C maps for both biological replicates in DN3 cells are shown at 5 kb resolution for a ∼1.2 Mb region, including the genes *Nfatc3* and *Cdh1*, to show reproducibility. Positions of genes are shown. C) Scatter plot for all normalized interaction scores from the two DN3 Capture Hi-C replicates. Spearman correlation coefficient (SCC) is shown on the graph. D) Pooled Capture Hi-C map for DP cells is shown at a ∼600kb region around the gene *Bcl6*. Dashed lines indicate a putative interaction between the *Bcl6* promoter and an upstream enhancer, identified by a peak of H3K27ac in the ChIP-seq track below. 4C-seq in DP cells using the *Bcl6* promoter as bait indicates sustained interactions over the broad upstream H3K27ac domain.

**Supplemental Figure 2. TADs are largely conserved, with cell type-specific differences**. A) Violin plots for distributions of insulation scores computed on pooled Hi-C datasets at 10 kb resolution, using a 150 kb (15 bin) window, on ESCs (green), DN3 (blue) and DP (red) cells. Whether analysis is restricted to the region targeted in the Capture Hi-C (top) or applied to the whole genome (bottom), ESCs have apparently more homogeneous insulation scores than thymocytes. B) Scatter plots comparing insulation scores from pooled Hi-C data at 10 kb resolution. Top: comparing ESC and DP cells using a 70 kb (7 bin) window; bottom: comparing DN3 and DP cells using a 150 kb (15 bin) window. SCC values are given on the graph. C) Plots of first two principal components for insulation scores computed on biological replicates of Capture Hi-C datasets at 5 kb resolution, using a 25 kb (5 bin) window (left) or a 75 kb (15 bin) window (right) for DN3 (blue), DP (red), untreated ESCs (green), and ESCs after either homozygous deletion of the major CTCF site at the *Bcl6* promoter (yellow) or CRISPRa ectopic induction of *Bcl6* (black) or *Nfatc3* (gray). D) Pairwise Venn diagrams for overlap of called TAD boundaries from the pooled Capture Hi-C data of ESCs (green), DN3 (blue) and DP (red) cells.

**Supplemental Figure 3. Cell type-specific TADs uncovered by conventional Hi-C**. A) Venn diagram for overlaps of called TAD boundaries from the pooled Hi-C data of ESCs (green), DN3 (blue) and DP (red) cells. B) Pairwise Venn diagrams for the overlaps, as in A). C) Pooled Hi-C maps shown at 10 kb resolution for a ∼2 Mb region around the *Runx1* gene in ESCs (top; green), DN3 (middle; blue) and DP (bottom; red) cells, just above color-coded plots showing the positions and scores of called TAD boundaries. Positions of genes are shown at the bottom of the plot. Red and blue bars denote the position of a thymocyte-specific domain formed around the *Runx1* gene, with a thymocyte-specific boundary denoted by an asterisk. D) Zoom on the thymocyte-specific domains shown in C) (top) and **Fig 2D** (bottom), showing the position of the genes and ChIP-seq tracks for CTCF in ESCs (green), DN3 (blue) and DP (red) cells. Red and blue bars denote the position of the thymocyte-specific domain and arrowheads above the plot denote the orientations of the relevant CTCF motifs. In both cases, convergent CTCF sites flank the new domain, and CTCF binding is enriched at one of these sites specifically in thymocytes.

**Supplemental Figure 4. The Bcl6 gene TAD border is broadened on DN3-to-DP transition**. Pooled Capture Hi-C plots at 5 kb resolution shown at a ∼600 kb region around the *Bcl6* gene in A) DN3; B) DP; C) untreated ESCs; and D) ESCs after CRISPRa to induce *Bcl6* expression. Positions of genes are given underneath. The white line shows the thyomocyte-specific promoter-enhancer contact point, which appears to demarcate the major TAD border in thymocytes.

**Supplemental Figure 5. Mild topological effects on deletion of the major CTCF site at the Bcl6 promoter in ESCs**. Capture Hi-C maps are shown at 5 kb resolution at a ∼600 kb region around the *Bcl6* gene in wild-type ESCs (top) and those with homozygous deletion of the major CTCF site at the *Bcl6* promoter (middle), as well as the differential map (bottom) comparing the two. Below are shown the positions of genes, the position of the deletion (red), the ChIP-seq track for CTCF in ESCs, and the plot of computed insulation scores at 2 kb with an 80 kb (40 bin) window for wild-type (green) and ΔCTCF (black) ESCs. The orientations of the main CTCF motifs are indicated by arrowheads. Dashed lines show sites of altered insulation: a loss of insulation at the deleted site, and a seemingly complementary gain of insulation at maintained downstream CTCF sites. Arrows on maps show where these insulation changes become apparent as stripes of relatively increased or decreased interactions.

**Supplemental Figure 6. Context-dependent transcriptional remodeling of TADs**. A-C) Capture Hi-C maps from which the differential maps in **Fig 4A** (A), **Fig 4B** (B), and **Fig 4C** (C) are derived are shown. Positions of genes are shown below, as well as ChIP-seq tracks for CTCF in DN3 (blue) and DP (red) cells (A-B), or ESCs (green; C). Differential CTCF binding does not seem to explain these topological changes. The rectangles represent domains around the activated gene specifically in DP (red; A), DN3 (blue; B) or induced ESCs (gray; C). D) Pooled Capture Hi-C maps are shown at 5 kb resolution for a ∼600 kb region around the gene *Il17rb* in DN3 (top) and DP (middle) cells, as well as the differential map (bottom) showing little difference between them. Below are shown the plot of insulation scores at 2 kb resolution with a window of 50 kb (25 bins) for DN3 (blue) and DP (red) cells, showing negligible differences and a maintained TAD border near the gene promoter, and ChIP-seq tracks for RNA polymerase II, showing much higher occupancy in DN3 (blue) cells than DP (red).

## Supplemental Tables

**Supplemental Table 1. Capture Hi-C strategy**. Overview of the genomic regions for which Capture Hi-C probes were designed, along with the relative expression differences between DN3 and DP cells for key genes within these regions.

**Supplemental Table 2. Overview of Hi-C and Capture Hi-C datasets presented in this study**.

**Supplemental Table 3. Reproducibility of Capture Hi-C**. Spearman correlation coefficients between all biological replicates performed in this study.

**Supplemental Table 4. Topological insulation similarities across cell type**. For pairwise combinations of the insulation scores, computed over different windows, their similarities are quantified with Spearman correlation coefficient, as well as their difference in distribution, as indicated by the p-values after two-tailed Kolgorov-Smirnov tests (KS p-val).

**Supplemental Table 5. Topological insulation similarities genome-wide across cell type**. As Table S4, but on the lower-resolution genome-wide Hi-C data. SCC values are given separately for the region targeted by Capture Hi-C and for the whole genome (“all”).

**Supplemental Table 6. Called TAD borders from Capture Hi-C experiments**. NA indicates where a TAD boundary was not called in that cell type.

**Supplemental Table 7. Called TAD borders from Hi-C experiments**. NA indicates where a TAD boundary was not called in that cell type.

**Supplemental Table 8. Sequences of gRNAs used in this study**.

**Supplemental Table 9. Sequences of qRT-PCR primers used in this study. Supplemental**

**Table 10. Oligonucleotides used in Capture Hi-C**.

**Supplemental Table 11. Sequences of 4C-seq primers used in this study**. Red sequence denotes Illumina sequencing adapters.

**Supplemental Table 12. Previously published GEO datasets used in this study**.

