## Supplemental Figures and Tables for "Context-dependent transcriptional remodeling of TADs during differentiation"

#### *Supplemental Tables*

**Supplemental Table 1. Capture Hi-C strategy.** Overview of the genomic regions for which Capture Hi-C probes were designed, along with the relative expression differences between DN3 and DP cells for key genes within these regions.

| **Gene** | **Expression change on DN3-to-DP transition** | **Number of probes** | **Captured region (mm10)** |
|---|---|---|---|
| *Bcl6* | 9.2-fold up | 1725 | chr16:23,659,574-24,258,391 |
| *Nfatc3* | 3.3-fold up | 3598* | chr8:105,833,089-106,979,086* |
| *Rag1* | 2.4-fold up | 1639 | chr2:101,357,817-101,956,585 |
| *Cdh1* | 6-fold down | 3598* | chr8:105,833,089-106,979,086* |
| *Il17rb* | 4.6-fold down | 1726 | chr14:29,680,447-30,279,827 |
| *Pla2g4a* | 4.6-fold down | 1578 | chr1:149,525,011-150,124,634 |
| *Cd3g* | n.s. | 1925 | chr9:44,666,642-45,265,845 |
| *Zap70* | n.s. | 1666 | chr1:36,456,898-37,055,359 |

* These two captured regions are fused as one larger region. n.s. = non-significant ($p > 0.05$).

**Supplemental Table 2. Overview of Hi-C and Capture Hi-C datasets presented in this study.**

| **Experiment** | **Number of reads** |
|---|---|
| DP Capture Hi-C, replicate 1 | 401,830,479 |
| DP Capture Hi-C, replicate 2 | 360,972,479 |
| DP DpnII Hi-C, replicate 1 | 148,715,287 |
| DP DpnII Hi-C, replicate 2 | 360,442,822 |
| DP HindIII Hi-C, replicate 1 | 176,274,361 |
| DP HindIII Hi-C, replicate 2 | 111,349,602 |
| DN3 Capture Hi-C, replicate 1 | 250,151,129 |
| DN3 Capture Hi-C, replicate 2 | 344,697,448 |
| DN3 DpnII Hi-C, replicate 1 | 160,220,248 |
| DN3 DpnII Hi-C, replicate 2 | 350,883,117 |
| DN3 HindIII Hi-C, replicate 1 | 140,836,381 |
| DN3 HindIII Hi-C, replicate 2 | 113,674,483 |
| ESC Capture Hi-C, replicate 1 | 342,156,642 |
| ESC Capture Hi-C, replicate 2 | 685,369,931 |
| ESC (CRISPRa *Bcl6*), replicate 1 | 306,192,847 |
| ESC (CRISPRa *Bcl6*), replicate 2 | 304,987,779 |
| ESC ΔCTCF (*Bcl6* promoter), replicate 1 | 328,152,115 |
| ES ΔCTCF (*Bcl6* promoter), replicate 2 | 360,340,711 |
| ESC (CRISPRa *Nfatc3*), replicate 1 | 307,926,726 |
| ESC (CRISPR *Nfatc3*), replicate 2 | 321,593,294 |

**Supplemental Table 3. Reproducibility of Capture Hi-C.** Spearman correlation coefficients between all biological replicates performed in this study.

| Experiment | Spearman correlation coefficient |
|---|---|
| DN3 | 0.96 |
| DP | 0.97 |
| ESC | 0.97 |
| ESC (CRISPRa *Bcl6*) | 0.96 |
| ESC ΔCTCF (*Bcl6* promoter) | 0.96 |
| ESC (CRISPRa *Nfatc3*) | 0.95 |

**Supplemental Table 4. Topological insulation similarities across cell type.** For pairwise combinations of the insulation scores, computed over different windows, their similarities are quantified with Spearman correlation coefficient, as well as their difference in distribution, as indicated by the p-values after two-tailed Kolgorov-Smirnov tests (KS p-val).

| Window | DN3 vs ESC | | DP vs ESC | | DN3 vs DP | |
|---|---|---|---|---|---|---|
| | **SCC** | **KS p-val** | **SCC** | **KS p-val** | **SCC** | **KS p-val** |
| 25 kb | 0.85 | $2.1x10^{-11}$ | 0.71 | $1.7x10^{-8}$ | 0.91 | 0.027 |
| 35 kb | 0.87 | $9.0x10^{-12}$ | 0.74 | $8.2x10^{-6}$ | 0.91 | 0.079 |
| 50 kb | 0.87 | $5.6x10^{-11}$ | 0.76 | $9.4x10^{-5}$ | 0.90 | 0.012 |
| 75 kb | 0.85 | $1.4x10^{-5}$ | 0.73 | 0.0041 | 0.91 | 0.12 |

**Supplemental Table 5. Topological insulation similarities genome-wide across cell type.** As Table S4, but on the lower-resolution genome-wide Hi-C data. SCC values are given separately for the region targeted by Capture Hi-C and for the whole genome ("all").

| Window | DN3 vs ESC | | | DP vs ESC | | | DN3 vs DP | | |
|---|---|---|---|---|---|---|---|---|---|
| | ***Capture region*** | | ***All*** | ***Capture region*** | | ***All*** | ***Capture region*** | | ***Genome*** |
| | ***SCC*** | ***KS pval*** | ***SCC*** | ***SCC*** | ***KS pval*** | ***SCC*** | ***SCC*** | ***KS pval*** | ***SCC*** |
| 70 kb | 0.65 | $1.5x10^{-9}$ | 0.49 | 0.42 | $<2.2x10^{-16}$ | 0.36 | 0.80 | 0.09 | 0.60 |
| 100 kb | 0.76 | $<2.2x10^{-16}$ | 0.57 | 0.51 | $<2.2x10^{-16}$ | 0.44 | 0.82 | 0.24 | 0.72 |
| 150 kb | 0.74 | $<2.2x10^{-16}$ | 0.65 | 0.51 | $<2.2x10^{-16}$ | 0.52 | 0.83 | 0.21 | 0.83 |

**Supplemental Table 6. Called TAD borders from Capture Hi-C experiments.** NA indicates where a TAD boundary was not called in that cell type.

Provided as separate Excel file.

**Supplemental Table 7. Called TAD borders from Hi-C experiments.** NA indicates where a TAD boundary was not called in that cell type.

Provided as separate Excel file.

**Supplemental Table 8. Sequences of gRNAs used in this study.**

| **Target** | **gRNA sequence** |
|---|---|
| *Bcl6* promoter (CRISPRa) | 5'-AGGGGAGGACTCGGTGGCAG-3' |
| | 5'-CCCGAGGCATTCTGCCGGCC-3' |
| | 5'-TGTTCCGGGCGGCGGTGCTG-3' |
| | 5'-CGTGACGGCGGCGGAGCGGG-3' |
| *Nfatc3* promoter (CRISPRa) | 5'-ATCGGGCGGAGCTCATGTCG-3' |
| | 5'-AGTCTCCAATTGGCCTACGT-3' |
| | 5'-TATCGCGTGAGTCCTCTGCG-3' |
| | 5'-CCAAGTTACGCCATCGAAGT-3' |
| *Bcl6* CTCF site (deletion) | 5'-GTGTAGAAGGCGATGCTAAC-3' |
| | 5'-TCACCGTTAATTCATACCGA-3' |

**Supplemental Table 9. Sequences of qRT-PCR primers used in this study.**

| **Gene** | **Sequence** |
|---|---|
| *Gapdh* | 5'-CATCACTGCCACCCAGAAGACTG-3' |
| | 5'-ATGCCAGTGAGCTTCCCGTTCAG-3' |
| *Bcl6* | 5'-CACGCGGTATTGCACCTT-3' |
| | 5'-CATCCACACAGGAGAGAAACC-3' |
| *Nfatc3* | 5'-CACCATCATTTCAGCTCCAA-3' |
| | 5'-GCACTCAAAGGGTTTAGGAC-3' |

**Supplemental Table 10. Oligonucleotides used in Capture Hi-C.**

Provided as separate Excel file.

**Supplemental Table 11. Sequences of 4C-seq primers used in this study.** Red sequence denotes Illumina sequencing adapters.

| **Name** | **Sequence** |
|---|---|
| *Bcl6* reading primer | 5'- AATGATACGGCGACCACCGAGATCTACACTCTTTCCCTACACGACGCTCTTCCGATCT CTTAAGGAGCCCACAGGAGTG-3' |
| *Bcl6* non-reading primer | 5'- CAAGCAGAAGACGGCATACGAGCTCTTCCGATCTGGGAGTCAAGGGATAAGACACA-3' |
| *Rag1* reading primer | 5'- AATGATACGGCGACCACCGAGATCTACACTCTTTCCCTACACGACGCTCTTCCGATCT AGGGACAAAACTTCTATTCATGATC-3' |
| *Rag1* non-reading primer | 5'- CAAGCAGAAGACGGCATACGAGCTCTTCCGATCTGGTCTCTCCCTTATATTCTTATCCTAA-3' |

**Supplemental Table 12. Previously published GEO datasets used in this study.**

| **Dataset** | **GEO accession number** | **Reference** |
|---|---|---|
| DN3 RNA-seq | GSE109125 | Yoshida et al., 2019 |
| DN3 CTCF ChIP-seq | GSE41743 | Shih et al., 2012 |
| DN3 RNA polymerase II ChIP-seq | GSE55635 | Pekowska et al., 2011 |

| | | |
|---|---|---|
| DN3 H3K27ac ChIP-seq | GSE80138 | Klein-Hessling et al., 2016 |
| DP RNA-seq | GSE109125 | Yoshida et al., 2019` |
| DP CTCF ChIP-seq | GSE41743 | Shih et al., 2012 |
| DP RNA polymerase II ChIP-seq | GSE29362 | Koch et al., 2011 |
| DP H3K27ac ChIP-seq | GSE63732 | Vanhille et al., 2015 |
| ESC CTCF ChIP-seq | GSE49847 | Yue et al., 2014 |
| ESC H3K27ac ChIP-seq | GSE66023 | Pradeepa et al., 2016 |

**Chahar et al. Fig S1**

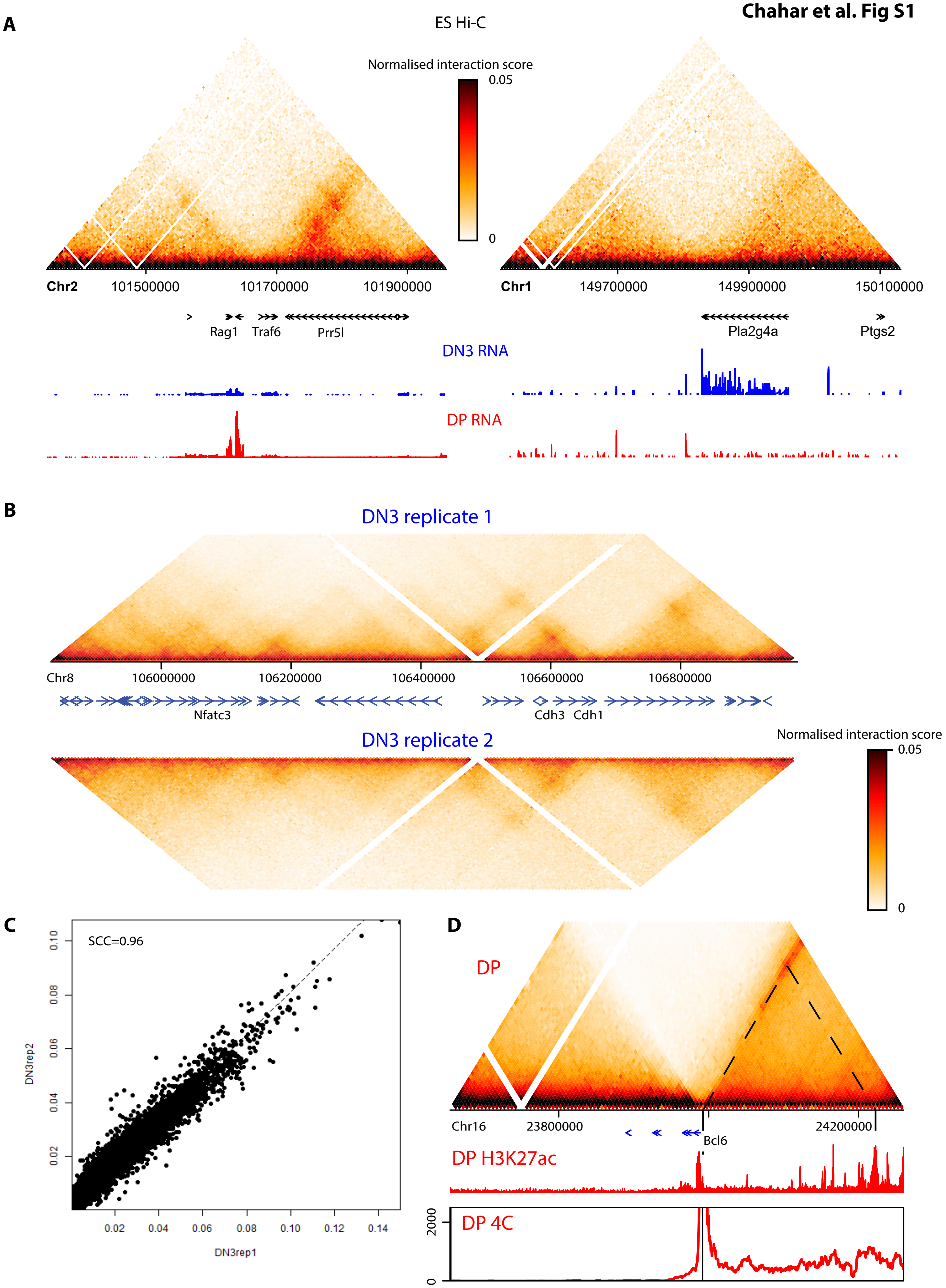

**Chahar et al. Fig S2**

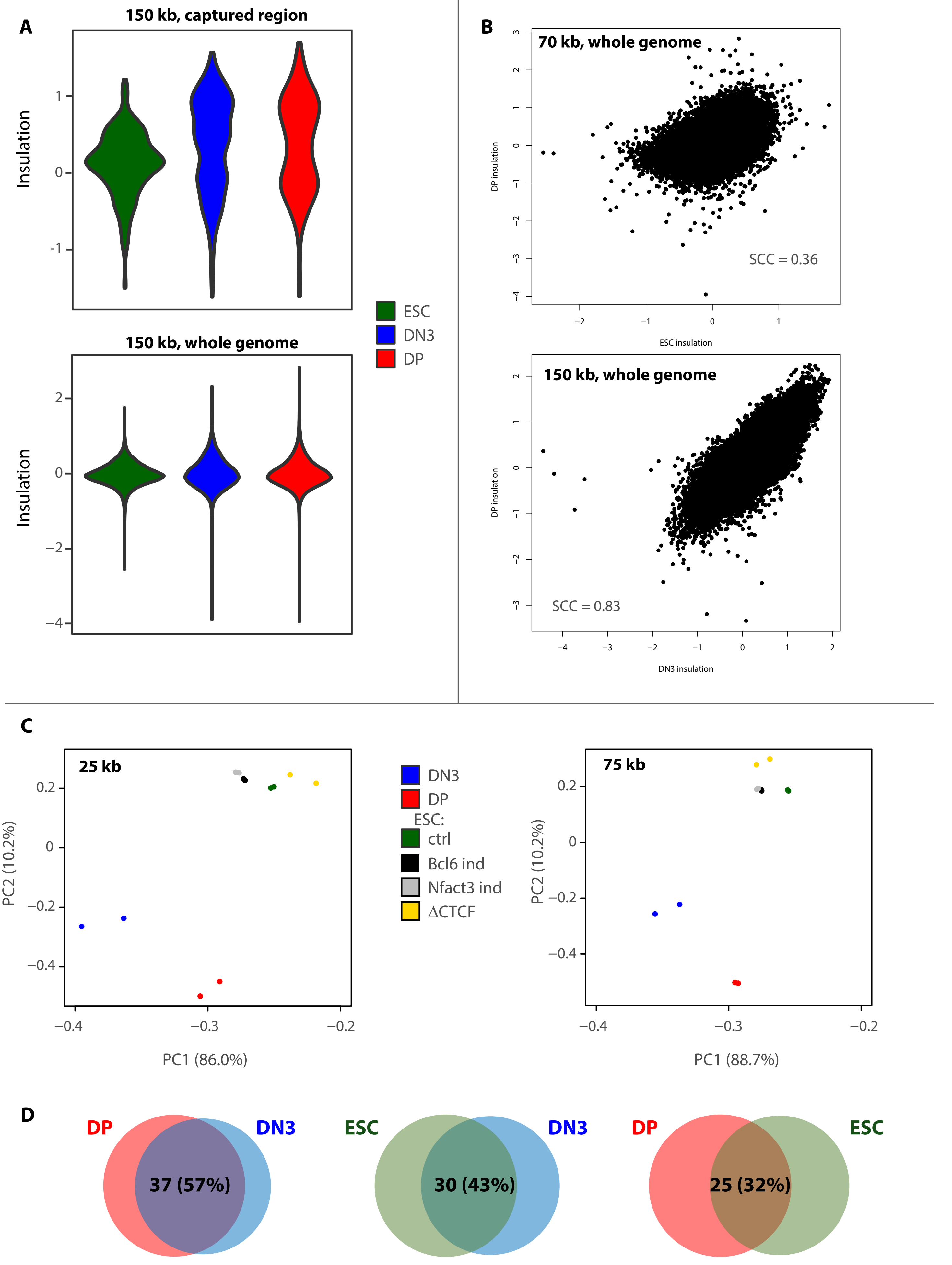

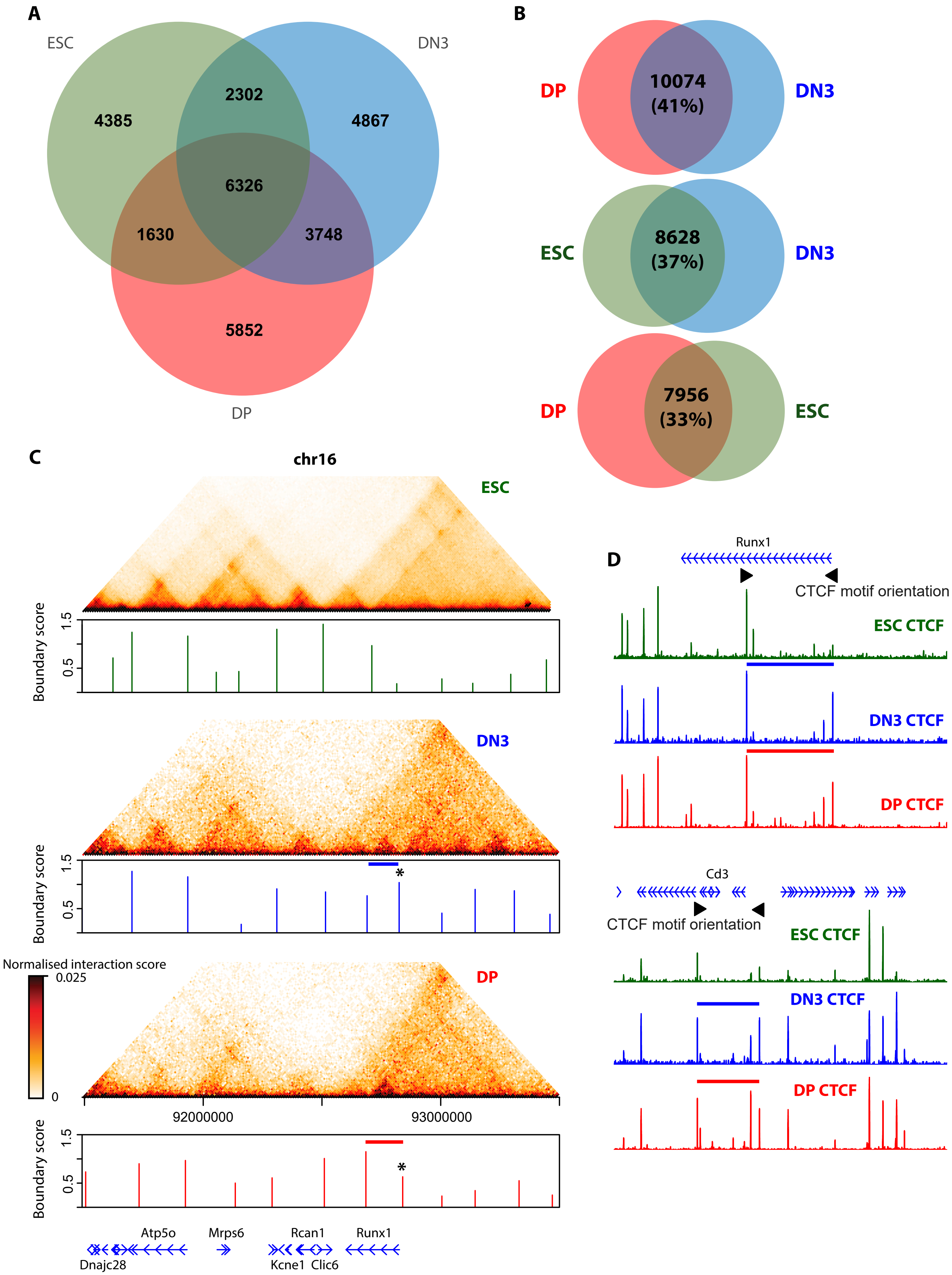
A
ESC
DN3
4385
2302
4867
6326
1630
3748
5852
DP
B
DP
10074
(41%)
DN3
ESC
8628
(37%)
DN3
DP
7956
(33%)
ESC
C
chr16
ESC
Boundary score
1.5
0.5
DN3
Boundary score
1.5
0.5
*
Normalised interaction score
0.025
0
DP
92000000
93000000
Boundary score
1.5
0.5
*
Atp5o
Mrps6
Rcan1
Runx1
Dnajc28
Kcne1
Clic6
D
Runx1
CTCF motif orientation
ESC CTCF
DN3 CTCF
DP CTCF
Cd3
CTCF motif orientation
ESC CTCF
DN3 CTCF
DP CTCF

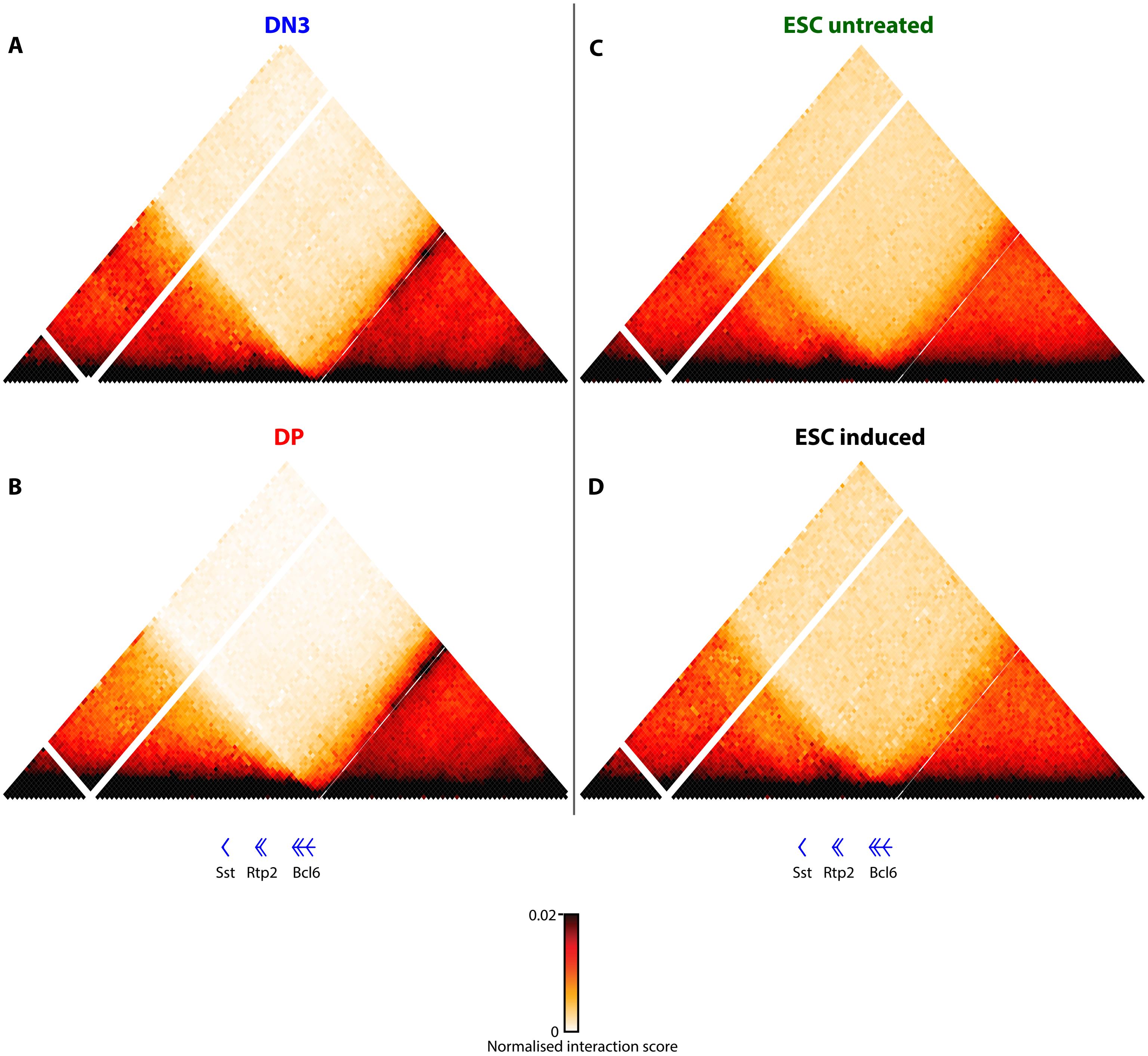
A
DN3
B
DP
C
ESC untreated
D
ESC induced
Sst
Rtp2
Bcl6
Sst
Rtp2
Bcl6
0.02
0
Normalised interaction score

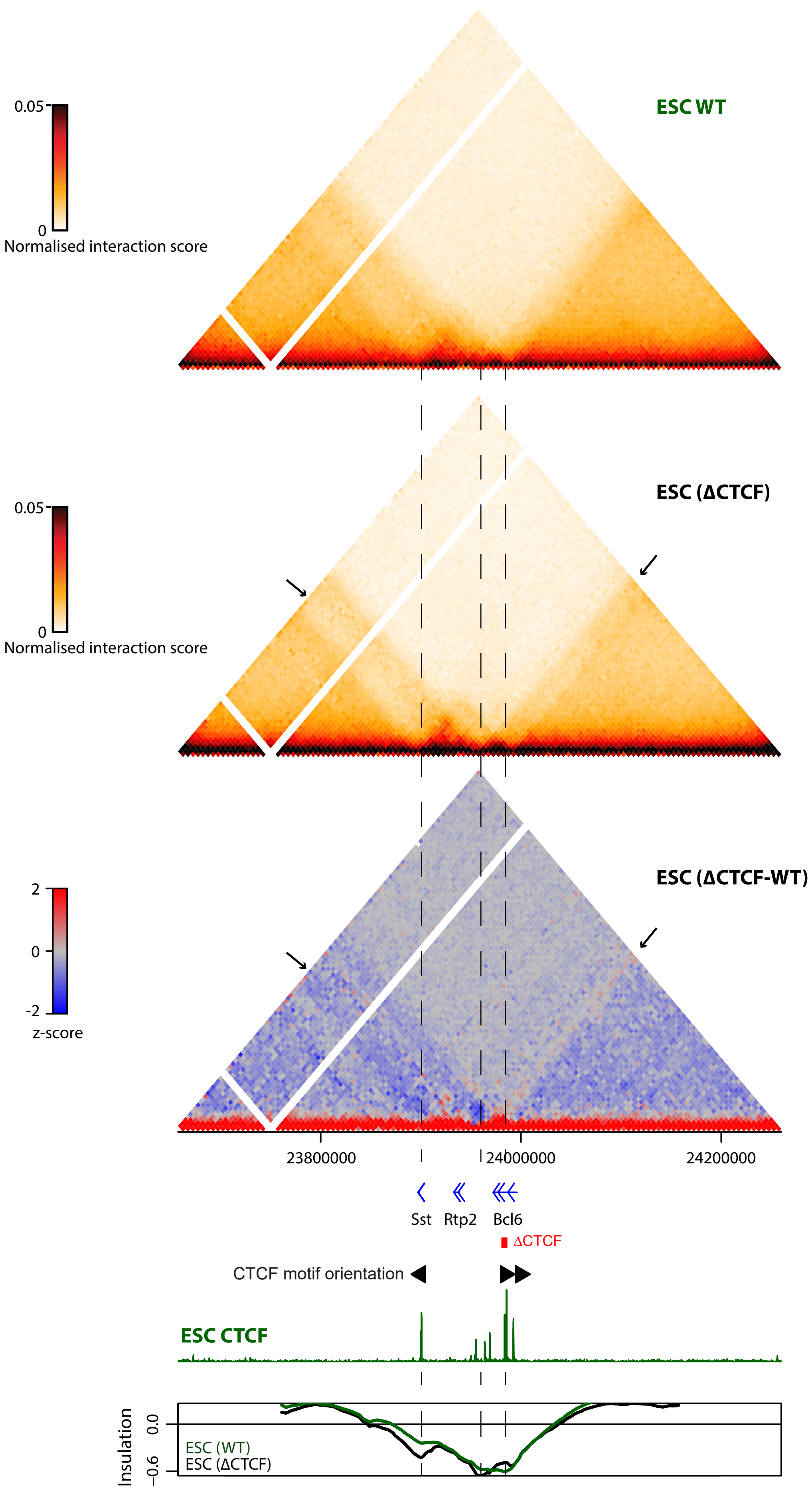
ESC WT
0.05
0
Normalised interaction score
ESC (ΔCTCF)
0.05
0
Normalised interaction score
ESC (ΔCTCF-WT)
2
0
-2
z-score
23800000
24000000
24200000
Sst
Rtp2
Bcl6
ΔCTCF
CTCF motif orientation
ESC CTCF
Insulation
0.0
−0.6
ESC (WT)
ESC (ΔCTCF)

### Chahar et al. Fig S6

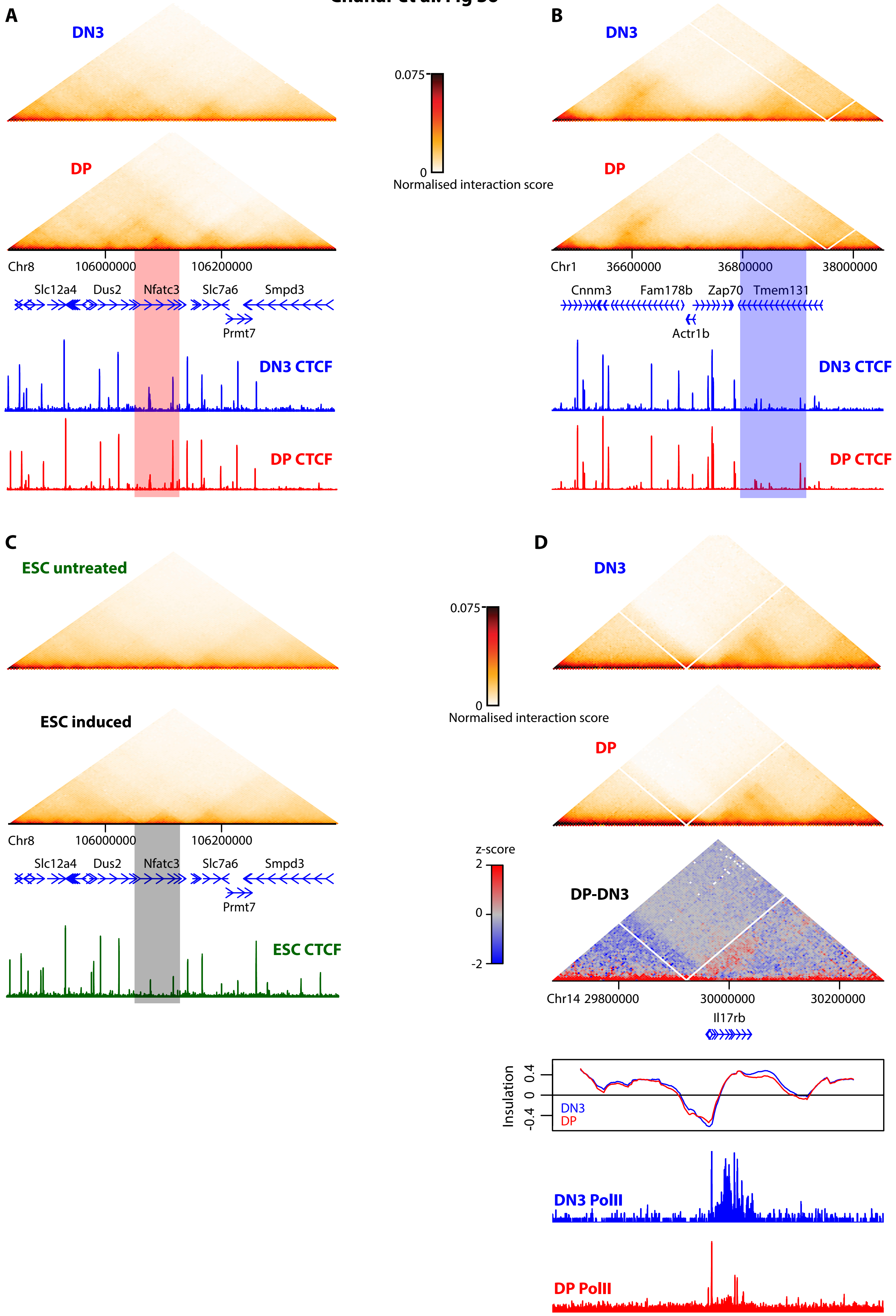
